# Chromosomal barcoding of *E. coli* populations reveals lineage diversity dynamics at high resolution

**DOI:** 10.1101/571505

**Authors:** Jesse Lerner, Michael Manhart, Weronika Jasinska, Louis Gauthier, Adrian W.R. Serohijos, Shimon Bershtein

## Abstract

Evolutionary dynamics in large asexual populations is strongly influenced by multiple competing beneficial lineages, most of which segregate at very low frequencies. However, technical barriers to tracking a large number of these rare lineages have so far prevented a detailed elucidation of evolutionary dynamics in large bacterial populations. Here, we overcome this hurdle by developing a chromosomal barcoding technique that allows simultaneous tracking of ∼450,000 distinct lineages in *E. coli.* We used this technique to gather insights into the evolutionary dynamics of large (>10^7^ cells) *E. coli* populations propagated for ∼420 generations in the presence of sub-inhibitory concentrations of common antibiotics. By deep sequencing the barcodes, we reconstructed trajectories of individual lineages at high frequency resolution (< 10^−5^). Using quantitative tools from ecology, we found that populations lost lineage diversity at distinct rates corresponding to their antibiotic regimen. Additionally, by quantifying the reproducibility of these dynamics across replicate populations, we found that some lineages had similar fates over independent experiments. Combined with an analysis of individual lineage trajectories, these results suggest how standing genetic variation and new mutations may contribute to adaptation to sub-inhibitory antibiotic levels. Altogether, our results demonstrate the power of high-resolution barcoding in studying the dynamics of bacterial evolution.

---

Advances in sequencing technologies have generated tremendous breakthroughs in identification of beneficial mutations arising in controlled laboratory evolution experiments, as well as mutations contributing to the emergence of anti-cancer or antibacterial drug resistance in the clinic^1–4^. Yet, experimental measurements of the *dynamics* of evolutionary processes remains a major challenge, particularly in large asexual populations, where multiple low-frequency small-effect mutations are known to spread simultaneously^5–8^. A quantitative description of evolutionary dynamics requires the ability to follow numerous individual lineages, most of which occur at extremely low frequency (10^−5^-10^−6^), and to do so in parallel and over multiple generations. Whole genome sequencing (WGS) techniques, although becoming routine and well-established, fall short of fulfilling this requirement, as they are usually unable to detect mutations at frequencies below ∼0.1%^9,10^. Various alternative solutions have been applied over the years to reconstruct population dynamics from trajectories of individual lineages at much higher resolution than accessed by WSG^11–13^. A particularly successful method that dramatically increases the frequency resolution of individual lineages is based on uniquely tagging chromosomes of individual cells with a genetic “barcode” that can be easily recovered by deep sequencing^14^. This approach was implemented in *S. cerevisiae*, where chromosomal insertion of ∼500,000 random barcodes using the *Cre-loxP* recombination system allowed a quantitative description of evolutionary dynamics of yeast populations (∼10^8^ cells)^5,15,15^. In bacteria, however, technical barriers have limited the number of uniquely-incorporated chromosomal barcodes to ∼100-400^17–19^. Such low levels of barcode diversity preclude us from answering important questions in bacterial evolution. For instance, highly-efficient chromosomal labeling is particularly relevant in characterizing the evolution of drug resistance in the presence of sub-inhibitory amounts of antibiotics, where the dynamics is driven by multiple mutations of low frequency and small fitness effects^20^.

Here, we present a method based on the Tn7 transposon to generate *E. coli* populations of > 10^7^ cells carrying 10^5^-10^6^ unique chromosomal barcodes. We demonstrate its implementation by quantifying the dynamics and patterns of lineage diversity loss in barcoded populations propagated in a month-long experiment (∼420 generations) in the presence of sub-inhibitory concentrations of two commonly applied antibiotics, chloramphenicol and trimethoprim. We found that different selection regimes elicited unique lineage diversity dynamics. By comparing the identity of individual barcodes in independent replicates evolving in parallel under identical conditions, we were also able to infer the relative contributions of pre-existing versus *de novo* mutations to the observed evolutionary dynamics. In general, stronger selection pressure generated faster loss of lineage diversity and more reproducible dynamics driven by standing genetic variation. In contrast, weaker selection pressure produced slower diversity loss and less reproducible dynamics due to a greater role of new mutations. In particular, ultra-low amounts of trimethoprim (0.01 μg/ml that amounts to 0.1% of minimal inhibitory concentration (MIC)) unexpectedly slowed the rate of lineage diversity loss even beyond conditions without any antibiotic, hinting the possibility that in this regime the antibiotic could be primarily functioning as a signaling molecule^21,22^.

## Results

### Highly efficient chromosomal barcoding of *E. coli* cells

Several robust genome-editing methods available for *E. coli* have long made this organism a flagship of genetic manipulations^23^. A high-resolution chromosomal labeling technique, however, has not been available, making it difficult to analyze the evolutionary dynamics of large bacterial populations segregating at low frequencies. To address this problem, we harnessed the well-established site-specific recombination machinery of the Tn7 transposon^24,25^. We placed the *tnsABCD* genes that encode the transposase biochemical machinery under the control of an arabinose-inducible pBAD promoter in a temperature-sensitive helper plasmid (**Fig. 1A**). The Tn7 arms (Tn7L, Tn7R) that target the genetic cargo at a neutral *attnTn7* attachment site were relocated to a suicide integration plasmid (**Fig. 1B**). We placed the barcode cassette carrying a 15-nucleotide long random sequence (the “barcode”) and the adjacent marker of selection between the Tn7 arms on the integration plasmid (**Fig. 1B,C**). To minimize the preparation of barcode libraries to two consecutive PCR reactions, we added sequences complementary to Illumina adapter primers flanking the barcode cassette (**Fig. 1C**). These sequences were used to both PCR amplify barcodes directly from cell cultures as well as anchor i5/i7 Illumina indexes to the amplified barcodes (Methods). Using this binary Tn7 transposon system, we integrated barcodes into a fixed location on the *E. coli* chromosome in two steps. First, we transformed cells with the Tn7 helper plasmid and pre-conditioned them by inducing the transposase machinery. Second, we transformed the pre-conditioned cells with the barcoded integration plasmid.

**Figure 1:**
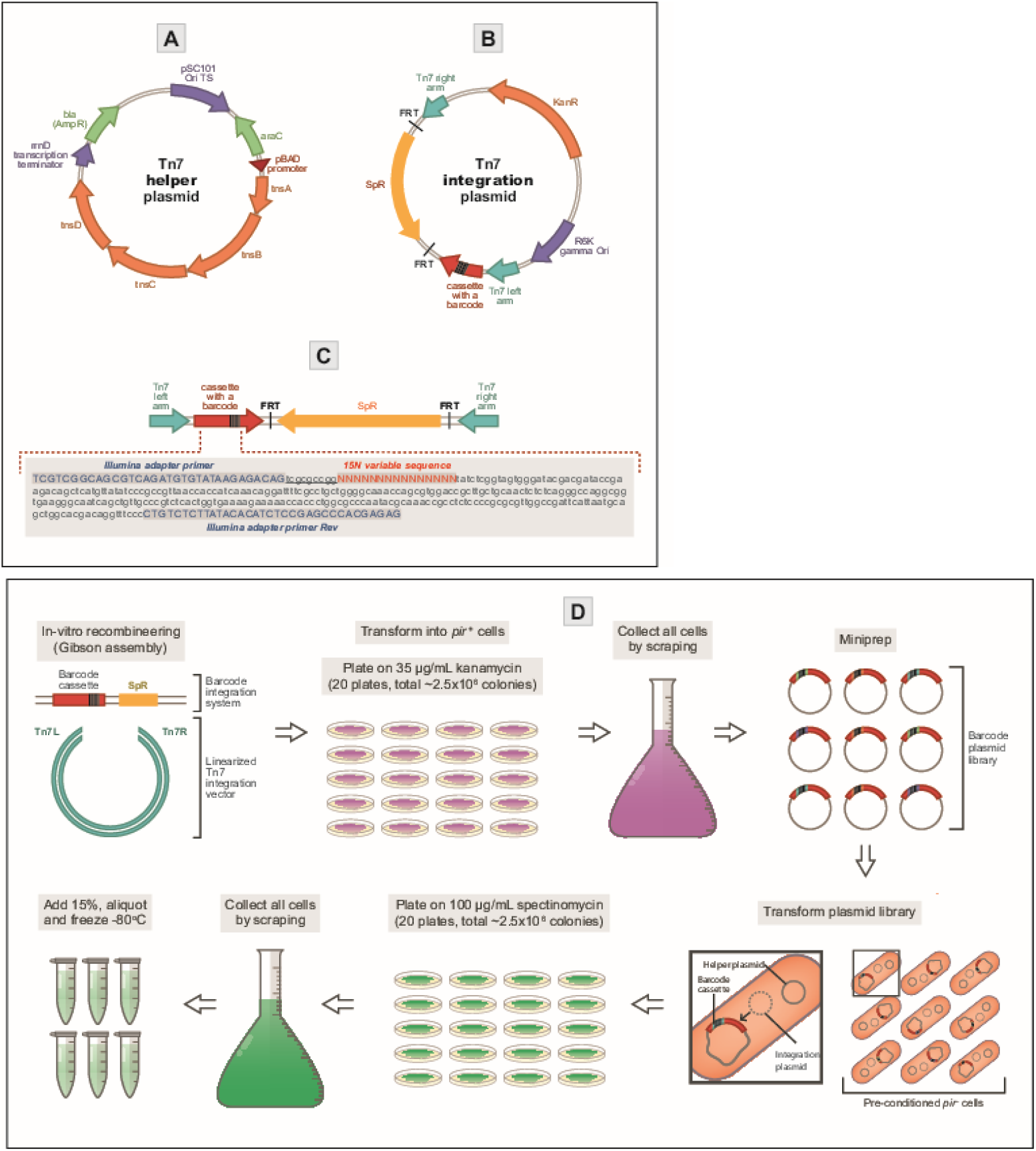
Barcoding *E. coli* cells with Tn7 transposon machinery. **(A)** Map of the helper plasmid expressing the Tn7 transposition machinery *(tnsABCD)* under the control of a pBAD promoter. Using a temperature-sensitive origin of replication (PSC101 Ori ts), we cure the plasmid by growing the strains at a non-permissive temperature after the chromosomal integration of barcodes is complete. **(B)** Map of a suicidal integration plasmid carrying the barcode cassette and spectinomycin resistance-conferring gene (SpR) nested between the left and right Tn7 arms. The SpR gene is flanked by FRT sites and can be later excised from the chromosome with Flp recombinase. The *pir+* dependent origin of replication (R6K gamma Ori) renders this plasmid suicidal in *pir-* cells. **(C)** A map of the segment undergoing chromosomal integration into the Tn7 attachment site *(attnTn7).* Inset: sequence of the barcode-carrying cassette. The 5’ and 3’ ends are flanked by sequences complementary to the Illumina adapter primers (upper case, blue) (see also Methods). The location of the 15 nt variable region (the barcode) is marked with uppercase red Ns. The barcode is placed upstream of a 9 nt stretch (underlined, lowercase). The barcode is the only variable part of the cassette. **(D)** Preparation of the barcoded plasmid and chromosomal libraries. We incorporate the cassettes carrying unique barcodes into Tn7 integration plasmids using Gibson assembly and then transform them into *pir+* cells. We achieve the chromosomal incorporation of the barcodes and curing of the Tn7 helper plasmids in a single plating step on spectinomycin at 37 °C (see also **Methods**).

We simultaneously selected for chromosomal barcode integration and removal of the helper plasmid by plating on selective media and incubating the plates at 37°C (**Fig. 1D, Methods**). Sequencing the ‘raw’ barcode library – as synthesized by the manufacturer and prior to incorporation of barcode cassettes into the integration plasmid (**Methods**) – revealed that the total number of unique barcodes was ∼1.3 × 10^6^ (**Table S1**). The ‘raw’ library had a fairly uniform distribution of frequencies: all barcodes but one had frequencies between 10^−7^ and 10^−5^ (**Fig. S1A,B**). The nucleotide composition of these barcodes was also very close to random, as quantified by the entropy of nucleotides per position (**Fig. S1C**). Incorporating the barcodes onto plasmids and then onto chromosomes reduced this diversity (∼8.4 × 10^5^ unique barcodes on plasmids and ∼4.5 × 10^5^ on chromosomes; **Table S1**). The process also introduced more redundancy into the distribution of frequencies, with some barcodes reaching frequencies of 10^−3^ (**Fig. S1A,B**). These increases in redundancy also led to a minor decrease in sequence entropy (**Fig. S1C**). However, the presence of a few barcodes with high initial frequencies did not appear to play a major role in the resulting lineage dynamics during evolution, as we show below.

### Laboratory evolution of the barcoded population

Bacteria are often exposed to antibiotic concentrations far below the minimal inhibitory concentration (MIC), both in natural environments and in patients receiving antimicrobial therapy^26,27^. Previous studies have shown that, compared to a lethal dosage, sub-MIC concentrations greatly expand the mutational space by allowing a large number of small-effect mutations to enter a population simultaneously^21^. The importance of low-frequency lineages in sub-MIC conditions make them a perfect setting to employ our barcoded population of *E. coli.* To this end, we chose two common antibiotics with distinct modes of actions: chloramphenicol (CMP), which inhibits protein synthesis via inactivation of peptidyl transferase activity of bacterial ribosome^28^; and trimethoprim (TMP), which functions as a competitive inhibitor of the essential protein dihydrofolate reductase^29^. It was demonstrated that sub-inhibitory concentrations of antibiotics as low as MIC/100 can still select for resistant mutants over the wild type^30^. Thus, to track the dynamics of adaptations at sub-MIC concentrations we chose antibiotic concentrations around MIC/100. To reduce the selection pressure even further, below the minimal selective concertation, we also chose ultra-sub-MIC concentrations (∼MIC/1000). To fine-tune the sub-MIC concentrations of both antibiotics to the conditions of the laboratory evolution experiment, we identified concentrations that reduced the total number of cells by no more than 30% by the end of a single propagation cycle (∼10 hours), comparatively to the untreated culture (**Methods**). These concentrations were 1 μg/mL CMP (6.25% MIC) and 0.1 μg/mL TMP (1% MIC) (**Fig. S2, Methods**). For ultra-sub-inhibitory regime, we chose 10% of the aforementioned concentrations. We then conducted laboratory evolution via serial passaging in the presence of CMP initially at 6.25% (‘low’) and 0.625% (‘ultra-low’) MIC, in TMP at 1% (‘low’) and 0.1% (‘ultra-low’) MIC, and in the absence of any antibiotics (**Fig. S3A,B**). We evolved 14 independent replicate populations in each of these five conditions (**Fig. S3C,D**). We diluted batch cultures (500 μL each) grown in 96 well plates by 1:100 every ∼6 generations (that is, passing twice daily; **Fig. S3E,F**) with a bottleneck population size of ∼3 × 10^7^ cells (**Fig. S3C,D**). To sustain the selection pressure along the evolutionary experiment (∼420 generations), we gradually increased CMP in the ‘low’ condition from 1 μg/mL to 2.8 μg/mL (**Fig. S3A**); and in the ‘low’ TMP condition, we increased the antibiotic from 0.1 μg/mL to 1.2 μg/mL (**Fig. S3B**). We kept the ‘ultra-low’ CMP environment constant at 0.1 μg/mL throughout the experiment (**Fig. S3A**), while ‘ultra-low’ TMP increased from 0.01 μg/mL to 0.1 μg/mL at generation ∼288 (**Fig. S3B**). As intended, the number of cells at the end of each passage remained roughly constant for each condition and along the entire evolutionary experiment (**Fig. S3C,D**).

In the experiment, we expected at least two forms of selective pressure, one due to the specific type and concentration of antibiotic, and another due to general growth conditions (growth medium, aeration, etc.). Additionally, there might also be fitness cost associated with the acquisition of drug resistance^31,32^. To analyze the effects exerted by these two selection forces, and a possible fitness cost, we measured the fitness of the barcoded populations at several time points during the experiment. Specifically, due to the limitation of the number of samples that can be loaded into a single NextSeq run, we chose a total of 12 randomly picked populations over the five conditions: three replicates for ‘low’ CMP, three replicates for ‘ultra-low’ CMP, two replicates for ‘low’ TMP, two replicates for ‘ultra-low’ TMP, and two replicates for ‘no drug’ condition. We measured fitness with respect to growth under antibiotics in units of IC50 (antibiotic concentration inhibiting 50% of growth) at the whole-population level (**Fig. S4, Methods**). These measurements revealed a moderate increase in antibiotic resistance over the experiment for ‘low’ CMP and TMP conditions (**Fig. S5A,B**). In contrast, both ‘ultra-low’ conditions produced no measurable improvement in IC50 (**Fig. S5A,B**).

Furthermore, the increases in drug resistance evolved in the ‘low’ conditions were accompanied by a fitness cost in the absence of antibiotics, measured by the growth rate (**Fig. S5C,D**). In particular, the population that evolved to resist the highest levels of CMP (‘low’ CMP replicate 1, **Fig. S5A**) showed the strongest fitness cost among the three replicate populations in the same condition (**Fig. S5C**). However, we observed no fitness cost in the populations evolved under ‘ultra-low’ CMP. Instead, the growth rate trajectories for these populations were generally similar to those for the populations that evolved in the absence of antibiotics (**Fig. S5C**). Surprisingly, in the populations evolved under ‘ultra-low’ TMP, the improvement in growth rate lagged behind the populations evolved under no antibiotics (**Fig. S5D**), suggesting that the rate of adaptation to growth conditions at 0. 1% MIC of TMP was diminished, despite the fact that no improvement in TMP IC50 was observed (**Fig. S5B**).

### Barcodes allow high-resolution monitoring of lineage trajectories

To elucidate the evolutionary dynamics of these populations at the level of individual lineages, we sequenced the barcodes of the same 12 populations at 16 time points. From the sequenced barcodes we assembled frequency trajectories for all ∼4.5 × 10^5^ initial lineages over the course of the experiment (**Figs. 2, S6**). These trajectories immediately raised several important qualitative insights. First, we saw that just one or two lineages dominate each population (> 85%) by the end of the experiment (see also **Table S2**). Second, some lineages rose to high frequency in several independent populations (lineage colors match across panels in **Fig. 2**). Third, we saw evidence of widespread clonal interference: some lineages that initially increased in frequency due to positive selection later decreased due to competition from fitter lineages. Furthermore, **Fig. 2** suggests that the rate of lineage diversity loss was reproducible between replicate populations in the same condition, while systematically distinct across different conditions. Although genetic drift alone can cause a drop in lineage diversity, this occurs on the time scale of 2*N* generations, where *N* is the effective population size^33^. Since the effective population size in our experiments was of order 10^7^ (**Fig. S3C,D**), neutral dynamics at the scale of the whole population was negligible on the time scale of the experiment. Therefore, we could assume that the dynamics of lineage diversity was dominated by selection.

**Figure 2:**
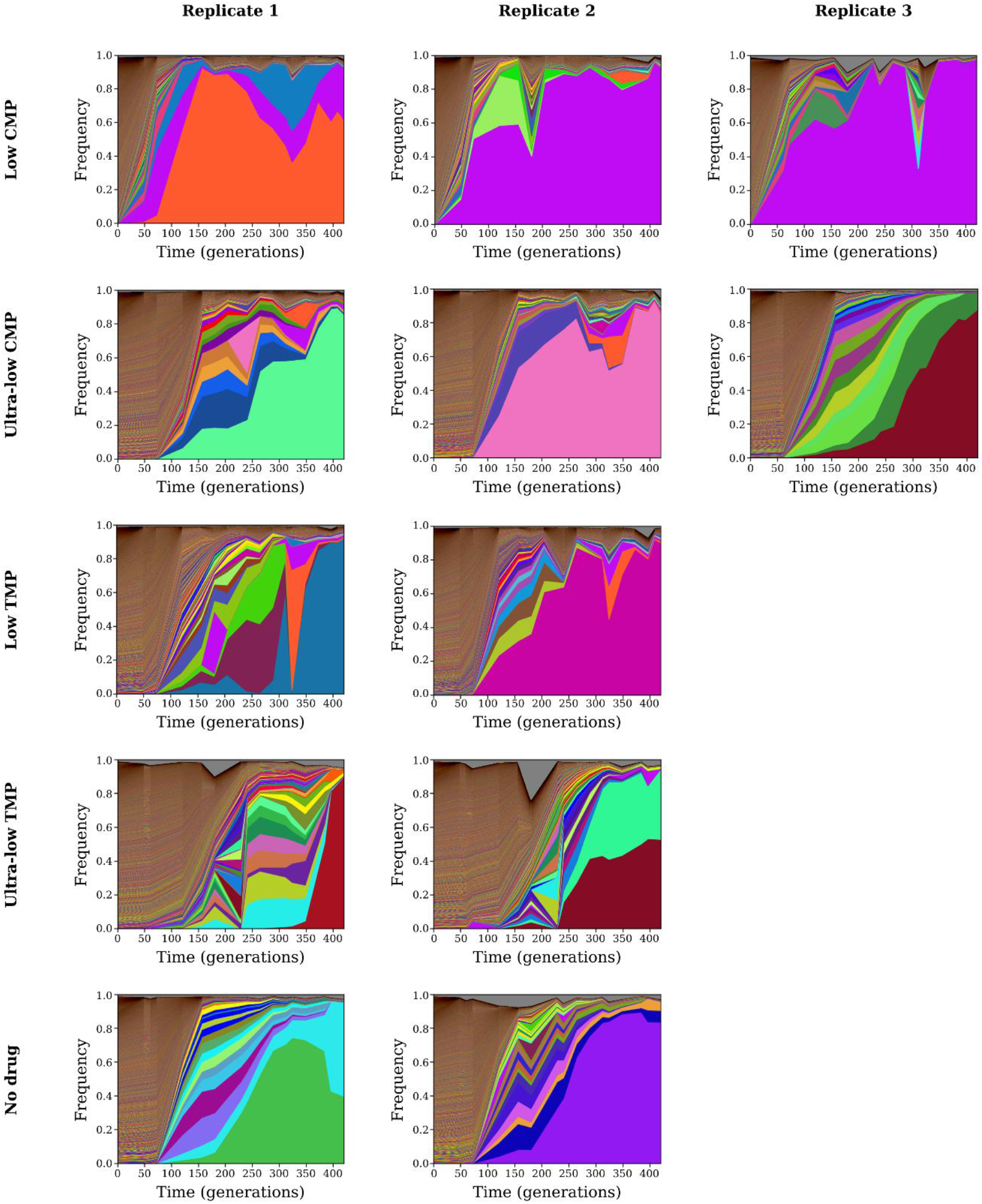
Dynamics of barcoded lineage frequencies over evolution experiment. Each panel shows the frequency trajectories for all barcoded lineages in a single population over time of the experiment. Each colored band corresponds to a unique lineage, with its vertical width indicating its frequency at a particular time point. The panels in each row correspond to a different antibiotic regimen, while each column corresponds to a different replicate. For the top 10 (according to average frequency) lineages in each population, we assign a unique color to each lineage that is consistent across panels (**Table S2**). We use random colors for all lower-frequency lineages, while gray represents the frequency of reads without identified barcodes.

### Ecological tools allow quantification of lineage diversity dynamics across conditions

To quantify the dynamics of lineage diversity, we adopted a measure of diversity widely used in ecological studies^34,35^:

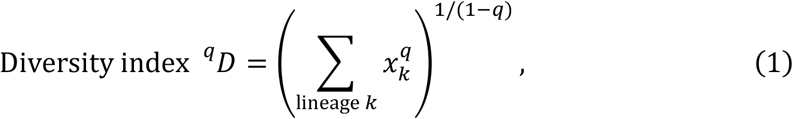

where *x_k_* is the frequency of the *k*^th^ barcoded lineage, and *q* is the “order” of the diversity index, which determines the sensitivity of diversity to abundant versus rare barcodes (**Methods**). In general, we can interpret the diversity index as the *effective* number of lineages present in the population. When *q* = 0, the diversity index simply counts the number of unique barcoded lineages, irrespective of their frequencies. This regime is equivalent to measuring diversity as “species richness” in ecological contexts^34^. When *q* = 1, the diversity index weighs all barcoded lineages by their frequency. This regime is equivalent to the exponential of the Shannon entropy of the frequencies. In the limit of *q* → ∞, the diversity index equals the reciprocal of the maximum lineage frequency, meaning that it depends only on the most abundant lineage and no others. Thus, by comparing the lineage diversity index across different *q* values, we can estimate the relative contributions of rare and abundant lineages to that diversity. Note, that if all lineages have equal frequencies, then the diversity index equals the actual number of lineages for any value of *q*. **Figure 3A** shows the dynamics of the lineage diversity index for each population over the time of the experiment, for three different values of *q*. At the beginning of the experiment, ^0^*D* ≈ 4.5 × 10^5^, since that is the total number of unique barcodes. However, the *effective* number of lineages, accounting for their unequal frequencies (**Fig. S1A,B**), is approximately 10-fold lower, ^1^*D* ≈ 4.6 × 10^4^. ‘Low’ CMP produced the fastest collapse of lineage diversity, extinguishing over 90% of unique barcodes in less than 50 generations. This behavior was displayed at all *q* values, indicating that rare and frequent barcodes contributed equally to these dynamics. In contrast, populations under ‘low’ TMP lost ^0^*D* diversity more rapidly compared to ‘ultra-low’ CMP and ‘no drug’ conditions, although it lost ^1^*D* and ^∞^*D* diversities at approximately the same rate; this indicates that the dynamics of abundant lineages were similar in these three conditions, but that low-frequency lineages disappeared more quickly in ‘low’ TMP. More surprising was the fact that populations under ‘ultra-low’ TMP lost diversity even more slowly than did populations under no antibiotics. Indeed, the diversity of populations under ‘ultra-low’ TMP maintained 40-50% of the initial effective diversity (*q* = 1) up to generation ∼120, whereas populations without antibiotics had only 1% of their initial diversity by that time point. This difference in the rates of lineage diversity deterioration between ‘ultra-low’ TMP and ‘no drug’ conditions is consistent with our fitness cost measurements, wherein the rate of adaptation under ‘ultra-low’ TMP lagged behind that of ‘no drug’ populations (**Fig. S5D**). Interestingly, *q* = 0 diversity under ‘ultra-low’ TMP is similar to the other populations by generation ∼250, but its diversity at larger *q* remains higher until the very end of the experiment. Towards the end of the experiment, we saw that the effective diversity of all populations is 1 or 2 lineages (*q* = 1), consistent with the observations from **Fig. 2**. However, we note that despite the drop in lineage diversity, there was still an ample number of surviving barcode lineages: each population retained a few thousand barcodes by the end of the experiment, as seen in the *q* = 0 diversity (**Fig. 3A**).

**Figure 3:**
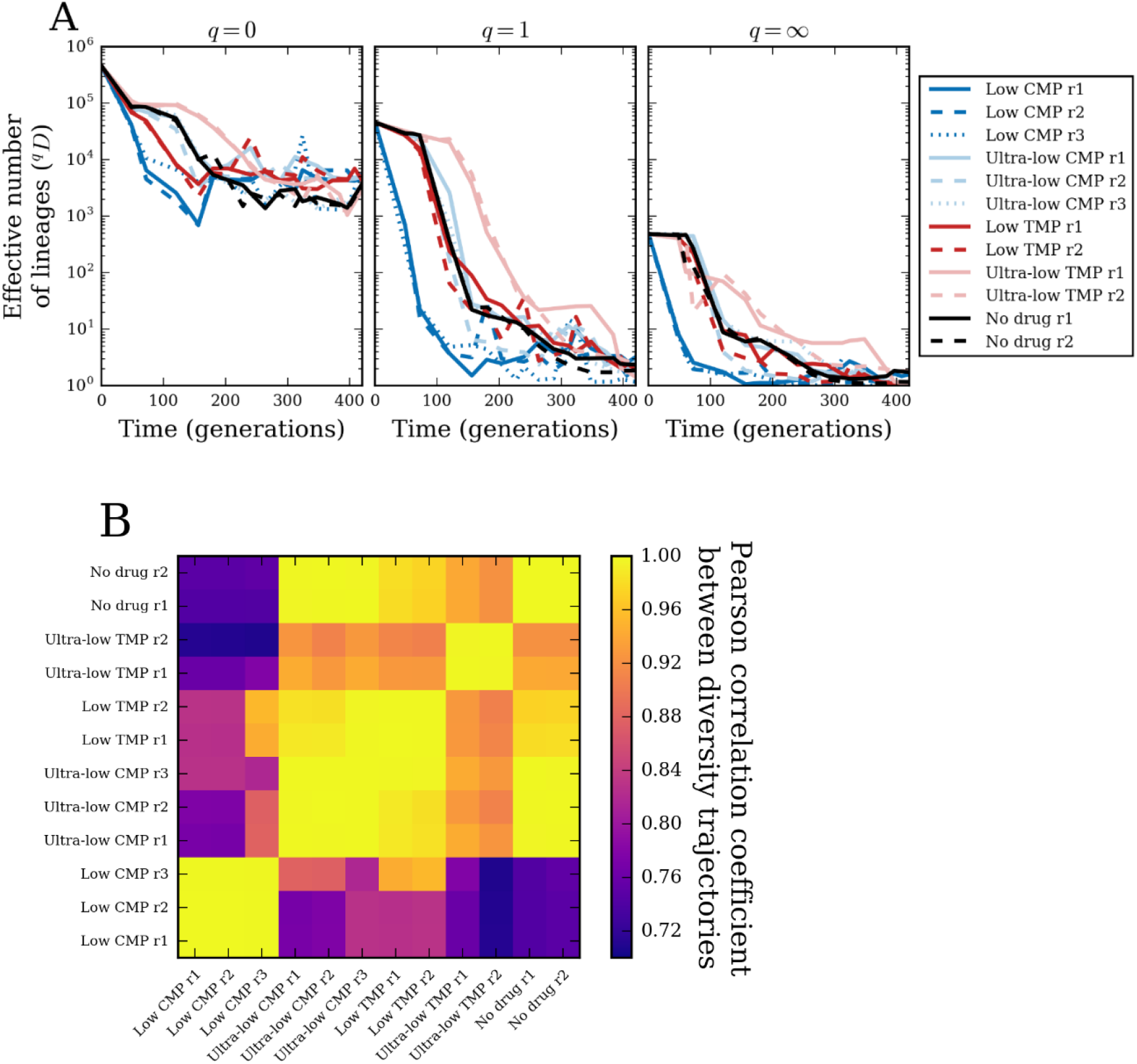
Dynamics of lineage diversity over time. We measure diversity of barcoded lineages using the index ^*q*^*D* (**Eq. 1, Methods**), where the parameter *q* controls the weight of low-versus high-frequency lineages: **(A)** for *q* = 0 (number of unique barcodes), *q* = 1 (exponential of Shannon entropy of lineage frequencies), and *q* = ∞ (reciprocal of the maximum lineage frequency). **(B)** Pearson correlation coefficient between diversity trajectories (^1^*D*) from all pairs of populations.

### Reproducibility of individual lineage dynamics

Not only did different antibiotic regimens produced distinct patterns of lineage diversity loss, but we also observed that these patterns are consistent across replicate populations. In **Fig. 3B** we show that the Pearson correlation coefficients between the *q* = 1 diversity trajectories from all pairs of populations. While all trajectories are somewhat correlated, since they all monotonically decrease, we saw stronger similarity among trajectories from the same condition. To further dissect whether individual lineages have similar fates across populations, we must quantify the similarity between lineage frequencies in different populations.

To compare the lineage composition of two or more populations at a single time point, we used a definition of diversity dissimilarity from ecology. Suppose we have *M* populations whose lineage compositions we want to compare. We first calculated the diversity index (**Eq. 1**) for all populations pooled together, ^*q*^*D*_pooled_ (**Methods**). We then calculated the diversity index for each population alone and determined the mean across all populations, ^*q*^*D*_mean_ (**Methods**). The ratio of these two quantities, shifted and rescaled, measures the dissimilarity among lineage compositions^34,35^:

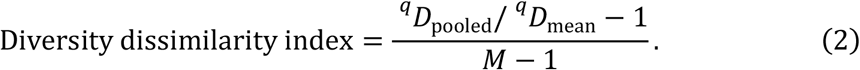

If the lineage compositions of all populations are identical, then the pooled population has diversity equal to the mean diversity, and the dissimilarity index equals zero. In contrast, if the lineage compositions of *M* populations have zero overlap, then the pooled population has diversity *M* times greater than that of the mean single population, and the dissimilarity index equals 1. As with the diversity index, the parameter *q* allows us to vary the importance of low and high-frequency lineages in the dissimilarity index. For *q* = 0, the dissimilarity index measures how many lineages multiple populations have in common, regardless of their frequencies, while for *q* → ∞, it compares only the highest-frequency lineages (**Methods**).

In **Fig. 4A** (left panel), we calculated the *q* = 1 dissimilarity between all pairs of populations at generation 120. At the beginning of the experiment, the populations were identical and so the dissimilarity between all pairs of populations was zero. By generation 120, we saw that many pairs of populations from different conditions have already diverged (dissimilarity close to the maximum value of 1), while pairs of populations from the same conditions remained more similar (except for ‘low’ TMP). Furthermore, we saw some similarity even between conditions: populations under the weakest antibiotic pressures (‘ultra-low’ CMP, ‘ultra-low’ TMP, and ‘no drug’) all maintained similarity between conditions comparable to their similarity between replicates. However, by the end of the experiment at generation 420 (**Fig. 4A**, right panel), we saw that most of this similarity between populations had disappeared. The main exception was ‘low’ CMP, where the three replicate populations maintained strong similarity. There was also a small amount of similarity between ‘low’ CMP, ‘ultra-low’ CMP (two out of three replicates), and ‘low’ TMP. In contrast, replicate populations under ‘ultra-low’ TMP and ‘no drug’ showed no similarity to each other by the end of the experiment. There was also strong similarity between replicate 3 in ‘ultra-low’ CMP and replicate 2 of ‘ultra-low’ TMP.

**Figure 4:**
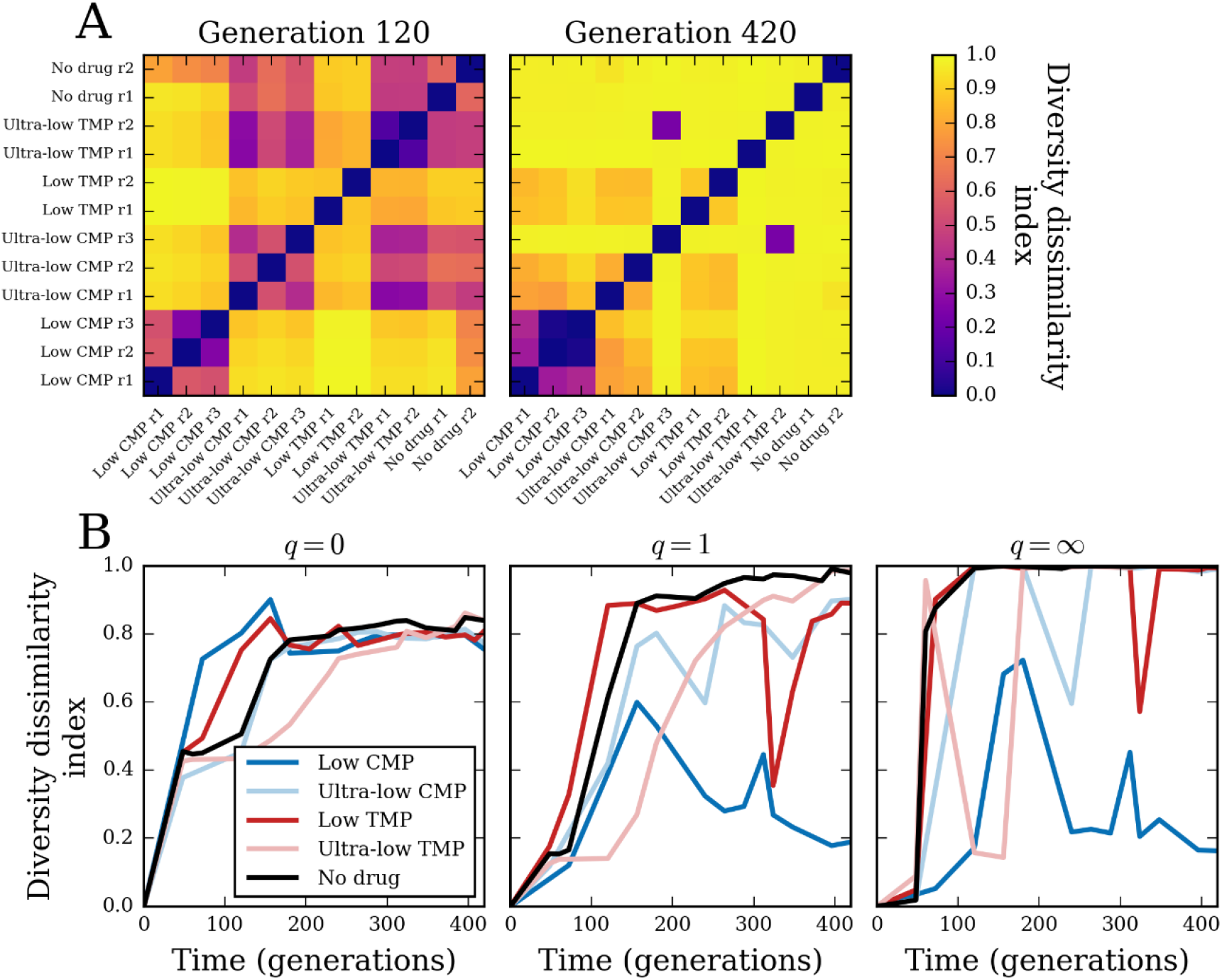
Dynamics of lineage dissimilarity among populations over time. **(A)** Diversity dissimilarity index for *q* = 1 (**Eq. 2, Methods**) between all pairs of populations at generation 120 (left) and the final time point at generation 420 (right). **(B)** Diversity dissimilarity index among all replicate populations in each condition, with *q* = 0, *q* = 1, and *q* = ∞.

We could further quantify the reproducibility of lineage dynamics by calculating diversity dissimilarity among all replicate populations in each condition over time (**Figs. 4B, S7**). For *q* = 0, we saw the dynamics of within-condition dissimilarity were similar to the dynamics of the diversity indices themselves in **Fig. 3A**. That is, populations under ‘low’ CMP diverged from each other most rapidly, followed by ‘low’ TMP, then concurrently by ‘ultra-low’ CMP and ‘no drug’, and, finally, by ‘ultra-low’ TMP diverging last. Interestingly, we saw all conditions settle at an intermediate amount of 0.8 dissimilarity by the end of the experiment; this value corresponds to having about 20% of their lineages in common (Methods). The dissimilarity index with *q* = 1, which accounts for heterogeneity in lineage frequencies, shows some differences with the *q* = 0 case that simply counts barcodes. With *q* = 1, ‘low’ CMP populations actually diverged more slowly from each other than did populations in the other conditions. Moreover, the ‘low’ CMP populations actually reached some maximum level of *q* = 1 dissimilarity around generation 150, and then began converging toward more similar lineage compositions. At the other extreme, ‘ultra-low’ TMP and ‘no drug’ populations reached maximum dissimilarity by the end of the experiment, while ‘low’ TMP and ‘ultra-low’ CMP populations had an intermediate value of dissimilarity. The dissimilarity index with *q* = ∞ shows a similar pattern of rise and fall for ‘low’ CMP populations; since this case depends only on the most frequent lineage, it implies that the high level of similarity between ‘low’ CMP populations by the end of the experiment was due to them sharing the same dominant lineage.

Altogether, this analysis shows that lineage dynamics under identical conditions are highly reproducible, even at the level of individual lineages. Specifically, it suggests that some lineages repeatedly rose to high frequency over multiple experiments. To demonstrate this more explicitly, in **Fig. 5A** we compared the frequencies of all barcoded lineages at the end of the experiment for two replicate populations in ‘low’ CMP, while in **Fig. 5B** we compared the trajectories for the top three lineages (ranked by average frequency over the experiment). Furthermore, in **Fig. 5C** we show the overlap of the top 10 barcodes between all three replicate populations in ‘low’ CMP: four of the top 10 in each population are shared among all replicates; with three more barcodes shared between two replicates (see also **Table S2**). Indeed, the most frequent lineage in replicates 2 and 3 is the exact same lineage (see purple trajectories in **Fig. 2**); this lineage is also the second-most frequent lineage in replicate 1. In contrast, **Fig. 5D,E,F** shows these same plots for the ‘no drug’ populations, which have little similarity among their most frequent lineages. In particular, they do not share any of their top 10 barcodes (**Fig. 5F, Table S2**). In **Figs. S8, S9** we show direct comparisons of lineage frequencies between all pairs of populations. The dominant lineages in each population do not appear to have started at unusually high frequencies (**Fig. S10**), suggesting another mechanism must explain their dominance.

**Figure 5:**
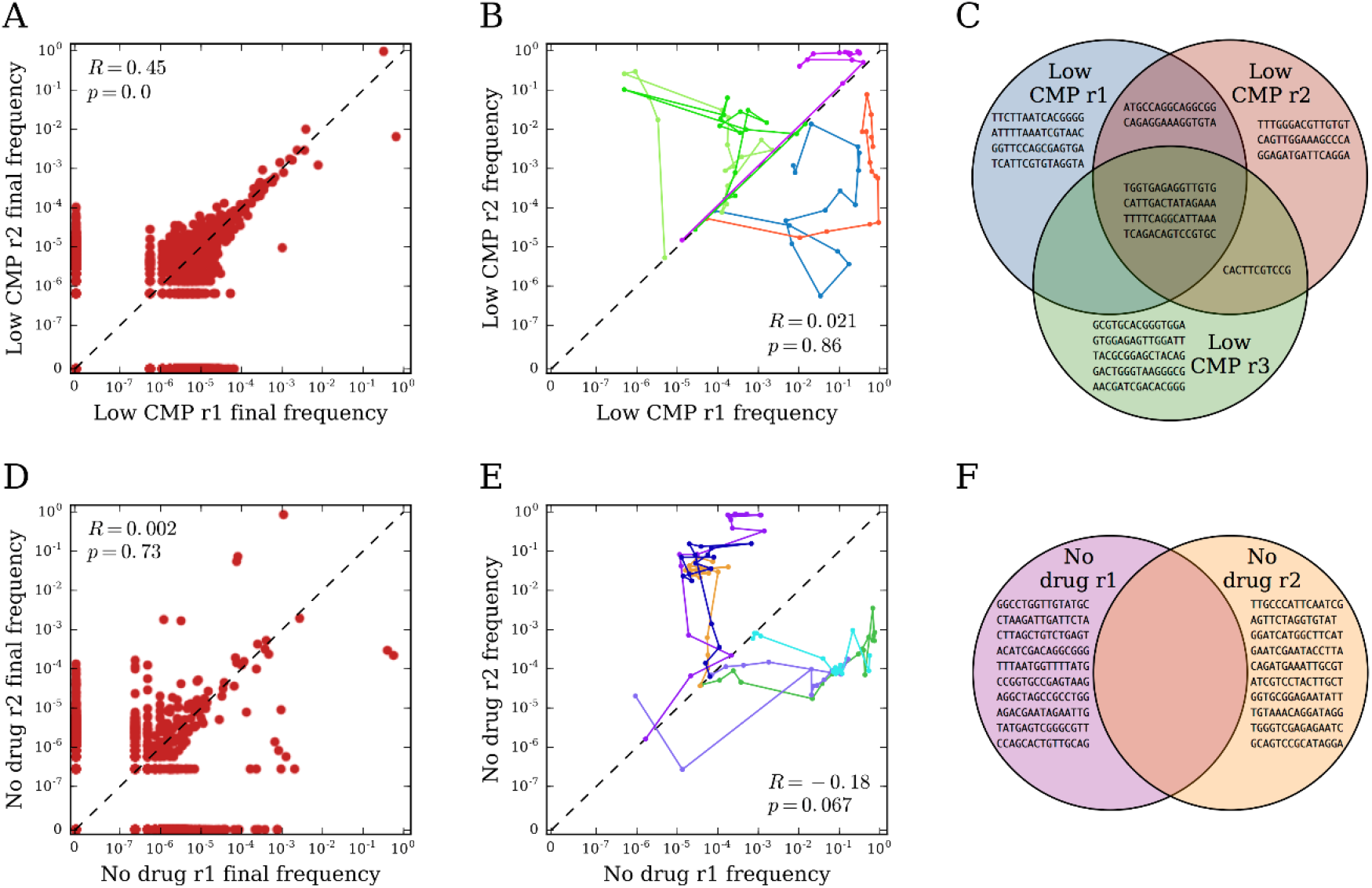
Repeatability of lineage dynamics. **(A)** Final frequencies of all barcoded lineages between two replicate populations in ‘low’ CMP. **(B)** Traces of lineage frequency over time for the union of the top three barcodes (by average frequency) in two replicate ‘low’ CMP populations. **(C)** Venn diagram of the top 10 barcodes (ranked by mean frequency over the experiment) in each replicate of ‘low’ CMP. **(D)** Same as (A) but for two replicate populations in ‘no drug’. **(E)** Same as (B) but for two replicate populations in ‘no drug’. In all panels, the dashed black lines mark the line of identity. **(F)** Same as (C) but for ‘no drug’ condition.

### Dynamics of individual lineages reveal clonal interference and the relative contribution of standing genetic variation versus and *de novo* mutations

One of the most salient features of individual lineages is that many of them appear to follow very similar trajectories (**Fig. S6**). To further elucidate the striking amount of similarity among some populations, we turned to the analysis of individual lineage trajectories in these populations. To that end, we performed hierarchical clustering of a subset of high-frequency lineage frequency trajectories in each population, based on the correlation coefficients between trajectories (**Figs. S11, S12**). We saw that the trajectories indeed formed well-defined clusters of distinct behaviors (**Fig. S13**); in particular, similar clusters appeared in replicate populations from the same condition, while more different clusters appeared in populations from different conditions. A few clustered trajectories also shared barcodes with very similar sequences, suggesting these lineages are actually the same (the distinct barcodes arising from sequencing errors) (**Fig. S14**). However, the fact that the vast majority of clustered trajectories involved unrelated barcode sequences suggests they are truly distinct lineages with highly correlated dynamics.

Arguably, the most interesting trajectory clusters are those that are non-monotonic with time, suggesting clonal interference. For example, **Fig. 6A,B,C** shows trajectory clusters from three populations in ‘low’ CMP with this property: these trajectories initially increased due to positive selection, but then later decreased as new mutations arose on other trajectories and outcompeted them. Every other population showed a similar cluster of trajectories, except those in ‘low’ TMP (**Fig. S13**); for example, **Fig. 6E,F** shows these trajectories in populations evolved without drug. The fact that these trajectories started increasing immediately suggests that beneficial mutations must have already been present on these lineages before the experiment. Indeed, the chromosomal barcoding process requires ∼30 generations of growth from the common ancestor. We surmised that during this time, random mutations began to accumulate in the population and, inevitably, were carried over to the evolution experiments. This also suggests an explanation for why some lineages rose to high frequency in multiple populations; these lineages likely carried beneficial mutations from the beginning, which allowed them to repeatedly dominate. We also observed a qualitatively different class of clonal interference trajectories, which did not start increasing immediately, but rose later in the experiment. For example, **Fig. 6G,H** shows these clusters in populations evolved with no drug. These trajectories appear to increase due to new mutations that arose during the experiment itself, rather than because of pre-existing mutations. We saw clusters of these trajectories in every population except those in ‘low’ CMP (**Fig. S13**). However, in replicate 1 of ‘low’ CMP, we do saw trajectories demonstrating both pre-existing beneficial mutations as well as new mutations, which initially rose due to the pre-existing mutations, then decreased from clonal interference, but then rose again due to the occurrence of a new mutation (**Fig. 6D**).

**Figure 6:**
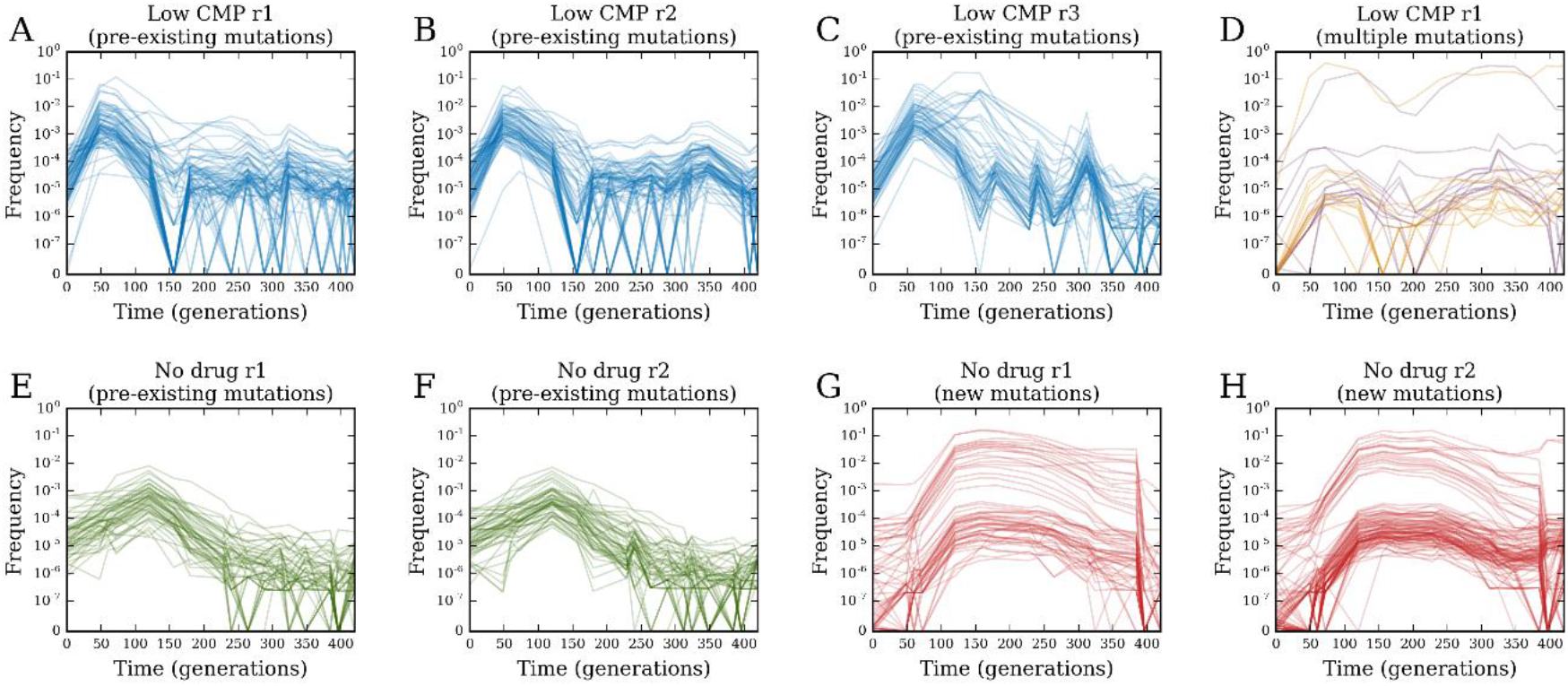
Distinct patterns of trajectories from pre-existing and new mutations. Clustered trajectories in **(A)** ‘low’ CMP replicate 1, **(B)** ‘low’ CMP replicate 2, and **(C)** ‘low’ CMP replicate 3 with putative pre-existing beneficial mutations. **(D)** Clustered trajectories in ‘low’ CMP replicate 1 with multiple beneficial mutations. Clustered trajectories in **(E)** ‘no drug’ replicate 1 and **(F)** ‘no drug’ replicate 2 with putative pre-existing beneficial mutations; clustered trajectories in **(G)** ‘no drug’ replicate 1 and (H) ‘no drug’ replicate 2 with putative new beneficial mutations.

## Discussion

Detailed understanding of evolutionary processes depends on our ability to follow individual lineages at a whole population level throughout multiple generations. Abundance of small-effect mutations in large populations makes it important to track lineages down to very low frequencies, ideally 10^−5^ – 10^−6^, in particular, during the initial rounds of evolution. However, methods for labeling lineages or whole-genome sequencing typically limit this resolution to orders of magnitude higher. Here we overcome this hurdle by developing a method that, for the first time, generates *E. coli* populations carrying 10^5^-10^6^ unique chromosomal barcodes. We tested the utility of the method by subjecting the barcoded population to serial passaging in presence of sub-inhibitory concentrations of common antibiotics. The relatively large size of the evolving populations (>10^7^) and the limited number of generations (∼420) have practically eradicated the contribution of drift to fixation of lineages in our experimental system, rendering selection the main force driving the loss of lineage diversity. Therefore, we can interpret the loss of lineage diversity as a proxy for the rate of adaptation in the population. We found that each condition prompted a different rate of adaptation (**Fig. 3A**), with the exception of ‘ultra-low’ CMP and ‘no drug’ conditions, which exhibited similar dynamics. Unexpectedly, ‘ultra-low’ TMP substantially reduced the rate of adaptation and lineage diversity loss, even compared to the rates observed for ‘ultra-low’ CMP and ‘no drug’ conditions. Since the growth dynamics of these populations indicate they experience a similar number of generations per passage as the other populations (**Fig. S3E,F**), we hypothesize that the observed delay in adaptation is due to a reduction in selection coefficients on beneficial mutations, compared to both the higher concentration of TMP (‘low’ TMP) and the condition without antibiotics. It was suggested that, at a low dosage, antibiotics might operate not as a weapon, but rather as signaling molecules that trigger transcriptional activation of multiple genes, including genes involved in the biosynthesis of amino acids, ribosomal proteins, purines, and pyrimidines^21^. If TMP, which is known to affect transcription^36^, indeed induces a new metabolic state in the bacterial cells at ultra-low concentration, it can potentially lead to a shift in the distribution of beneficial mutations. Namely, the fitness advantage of mutations under growth in the absence of antibiotics can decrease in the presence of 10 ng/ml of TMP, which is the ultra-sub-MIC concentration used in our experiment. It is important to note that we could not have easily detected a delayed adaptation in populations subjected to ‘ultra-low’ TMP, had we not used the barcode sequencing approach. Investigating the mechanism underlying the delayed adaptation at ‘ultra-low’ TMP is the subject of another work in the immediate future.

One of the important problems in evolutionary biology is robustly quantifying the reproducibility of evolutionary processes^37,38^. The deterministic nature of adaptation at near-lethal drug concentrations has been demonstrated at the level of a handful of strongly-beneficial mutations that have repeatedly accumulated in almost pre-determined order in replicate populations^39^. However, quantifying the reproducibility of evolutionary dynamics driven by many small-effect mutations at the whole-population level has remained elusive in bacteria. Our chromosomal barcoding system in *E. coli* allowed us to directly address this problem by quantitatively comparing lineage dynamics across independent replicate populations subjected to identical antibiotic regimes. We found that the rate of lineage diversity loss was highly reproducible for each selection condition (**Fig. 3A**), indicating that lineage diversity analysis provides a robust way to quantify the reproducibility of evolution at a whole-population level. Curiously, the diversity dynamics were reproducible regardless of the rate of adaptation, For example, both ‘low’ CMP condition, which induced the fastest rate of adaptation, and ‘ultra-low’ TMP condition, which induced the slowest adaptation dynamics, exhibited high reproducibility between evolutionary replicates. This observation implies a surprising deterministic dynamics at the level of lineage diversity, despite differences in the distributions of fitness effects across conditions in our experiment.

The dynamics were furthermore deterministic at the level of individual lineages in some cases, with a few lineages rising to high frequency in multiple independently evolving populations. Populations can adapt to new environments using two sources of beneficial mutations: pre-existing mutations (i.e., standing genetic variation) and new mutations^40^. These two mechanisms can have different effects on the rates and outcomes of evolution, yet delineating the evolutionary dynamics with respect to the relative contributions of preexisting and new mutations is nontrivial. Competition experiments in presence of sub-MIC amounts of antibiotics demonstrated that pre-existing mutations can be enriched for by sub-MIC selection^30,41^. Sub-MIC selection was also shown to increase the frequency of new resistance-conferring mutations in evolving populations^30,42^. However, neither the relative contribution of standing genetic variation versus novel mutations in a single population under sub-MIC selection, nor the evolutionary dynamics resulting from the interplay between these two mutational sources have been demonstrated. Here, we developed a quantitative approach to address this problem. By applying a measure of lineage composition dissimilarity from ecology^34,35^, we compared the change in frequency of identical lineages between independent replicates subjected to identical conditions over the evolutionary time course. These results suggest that lineages present at very low frequencies at the initial population, but nonetheless independently reaching high frequency in the replicate populations, are repetitively selected for because they carry preexisting beneficial mutations. Thus, the extent of similarity in lineage composition between evolutionary replicates at a particular time point quantitatively reports on the contribution of standing genetic variation to the observed dynamics. Expectedly, the rapid adaptation under ‘low’ CMP was accompanied by the lowest dissimilarity index among selection regimes (**Fig. 4**), indicating a heavy contribution of standing genetic variation. Conversely, very little indication of the contribution of pre-existing variation can be seen for ‘ultra-low’ TMP and ‘no drug’ conditions (**Fig. 4**), indicating that the evolutionary dynamics of the latter was driven mostly by newly acquired mutations. Further support for the importance of pre-existing beneficial mutations comes from our analysis of individual trajectories. Lineages increasing in frequency due to a newly-acquired beneficial mutation show a finite establishment time (time required for a lineage prior to committing to a deterministic growth after acquisition of a mutation). In contrast, lineages carrying pre-existing mutations increase immediately upon applying selection pressure, with zero establishment time. Indeed, among all the identified individual lineages exhibiting clonal interference trajectories (**Fig. S13**), no establishment time can be seen under ‘low’ CMP conditions, while finite establishment times can be detected in all other conditions.

Overall, our results demonstrate significant evolutionary insights gleaned from high-resolution lineage tracking using chromosomal barcodes in *E. coli.* Our experimental barcoding protocol based on the Tn7 transposon machinery is straightforward to implement and can readily be reproduced in a variety of systems. We have furthermore shown how to obtain a robust quantitative analysis of the resulting lineage data using ecological diversity indices. Altogether, we envision that this tool will find wide applicability in addressing diverse questions in bacterial population and evolutionary dynamics.

## Methods

### Design of the chromosomal barcode integration system in *E. coli*

The design of the chromosomal integration system of the barcode library is based on the pGRG25 plasmid, which carries all the components of the Tn7 transposon site-specific recombination machinery^25,43,43^. We separated the Tn7 transposase machinery (tnsABCDE) from the Tn7 arms (Tn7L, Tn7R) onto two independent plasmids: the temperature-sensitive *helper* plasmid (pSC101 temperature-sensitive origin of replication) carrying the *tnsABCD* genes under the control of an arabinose-inducible pBAD promoter (**Fig. 1A**), and the suicide *integration* plasmid (R6K gamma *pir+* dependent origin of replication) with the Tn7 arms flanking the barcode-carrying cassette and spectinomycin resistance gene (**Fig. 1B,C**). The separation of the recombination machinery *tnsABCD* from the integration segment flanked by the Tn7 arms achieves two major goals. First, the introduction of the helper plasmid into the cells prior to the integration plasmid allows us to precondition the cells for chromosomal recombination by inducing the expression of the transposase complex, which is expected to increase the integration efficiency. Second, the transient nature of the integration plasmid will eliminate any possibility of barcodes lingering outside of their designated chromosomal location due to unsuccessfully cured plasmids. This is especially important for barcoding temperature-sensitive strains that cannot be grown at non-permissive temperatures. The design of the barcode carrying cassette (**Fig. 1C**) is aimed at minimizing the number of library preparation steps required for Illumina deep-sequencing technology. The only variable region in the cassette is the barcode sequence of 15 random nucleotides. Thus, the barcode library has a theoretical diversity of 4^15^ (∼10^9^) unique barcodes. The barcodes are asymmetrically located 9 nt downstream of the 5’ end of the cassette. This short stretch of 9 nt is sufficient to locate the barcodes in the raw sequencing data files and to reduce the sequence redundancy in the Illumina flow cell. The cassette is flanked by the sequences complementary to the Illumina adapter primers used to anchor library specific indexes recognized by the Illumina sequencing platforms (MiSeq, HiSeq, or Nextseq) (**Fig. 1C**). The Integrated DNA Technology (IDT) gBlocks service synthesized the barcode cassettes (allowing the incorporation of 15 consecutive and variable nucleotides), which we then cloned into the Tn7 integration plasmid (**Fig. 1B**). We characterized the resulting library of barcodes by deep sequencing prior to integration into the genome (**Fig. S1**).

### Design of the barcode-carrying cassette

The barcodes comprise 15 random nucleotides, which we placed 9 nt downstream of the 5’ end of the 288-nt long cassette:

g<u>tcgcgccgg</u>NNNNNNNNNNNNNNNtatctcggtagtgggatacgacgataccgaagaca gctcatgttatatcccgccgttaaccaccatcaaacaggattttcgcctgctggggcaaaccagcgtggaccgcttgctgcaactctctcagggccaggcggtgaagggcaatcagctgttgcccgtctcactggtgaaaagaaaaaccaccctggcgcccaatacgcaaaccgcctctccccgcgcgttggccgattcattaatgcagctggcacgacaggtttccc

We placed the barcode-carrying cassette between sequences complementary to Illumina adaptor primers (forward overhang: 5’ TCGTCGGCAGCGTCAGATGTGTATAAGAGACAG; reverse overhang: 5’ GTCTCGTGGGCTCGGAGATGTGTATAAGAGACAG) and, finally, flanked by sequences complementary to the integration site in the Tn7 integration plasmid:

<u>gatatcggatcctagtaagccacgttttaattaatcagatccctcaatag</u>ccacaacaac tggcgggcaaacagtcgttgctgattggtcgtcggcagcgtcagatgtgtataagagaca g<u>tcgcgccgg</u>NNNNNNNNNNNNNNNtatctcggtagtgggatacgacgataccgaagaca gctcatgttatatcccgccgttaaccaccatcaaacaggattttcgcctgctggggcaaa ccagcgtggaccgcttgctgcaactctctcagggccaggcggtgaagggcaatcagctgt tgcccgtctcactggtgaaaagaaaaaccaccctggcgcccaatacgcaaaccgcctctc cccgcgcgttggccgattcattaatgcagctggcacgacaggtttcccctgtctcttata cacatctccgagcccacgagac<u>gccactcgagttatttgccgactaccttggtgatctcgcctttcacgtag</u>

Integrated DNA Technology (IDT) (https://www.idtdna.com/pages/products/genes-and-gene-fragments/gblocks-gene-fragments) synthesized the resulting 492 nt-long sequence as double-stranded gBlock with randomly mixed bases (at the 15 consecutive positions indicated by N).

### Generation of plasmid barcode library

We digested the empty Tn7 integration plasmids with NotI, purified them by ethanol precipitation, and then mixed the plasmids with gBlocks in a 1:3 molar ratio in the presence of NEBbuilder HiFi DNA (New England Biolabs) assembly mix according to the manufacturer’s instructions. We used a control without NEBbuilder HiFi DNA assembly mix to determine the background level (Tn7 integration plasmid lacking the cassette). We concentrated the reactions by ethanol precipitation and transformed them into 100 μL of TransforMax™ EC100D™ *pir+* cells (Lucigen). We resuspended the transformed cells in 1 mL of SOC medium, regenerated them for 1 hour at 37 °C with shaking followed by overnight incubation on the bench, and then plated the cells on 35 μg/ml kanamycin LB agar plates (100 μL of transformed cells per plate). We plated several microliters of diluted cultures separately to determine the number of colony-forming cells. We incubated plates overnight at 37 °C. Cells transformed with DNA but without the assembly mix produced no colonies. Transformation of DNA with the assembly mix generated a total of ∼2.4 × 10^6^ colonies (**Fig. 1D**). We scraped all the colonies from the plates, pooled them together, and thoroughly mixed them. Finally, we extracted plasmids from the pooled cells with a Qiagen midi kit.

### Chromosomal integration of the barcode library into *E. coli* cells

This is a two-step process, the first step being the transformation of the recipient *E. coli* MG1655 cells with the Tn7 helper plasmid and induction of the transposase integration machinery. The second step is the transformation of the Tn7 integration plasmid library, which will integrate the barcodes into the chromosome (**Fig. 1D**). We grew cells transformed with the Tn7 helper plasmid overnight until saturation in LB supplemented with 100 μg/mL ampicillin at 30 °C. We then diluted these cells 1:100 and grew them under the same condition for 45 min. We added 0.2% arabinose and diluted cells to OD600nm of 0.5. We then harvested the cells, washed them 3 times with ice-cold water, and transformed them with the Tn7 integration plasmid library using electroporation. We resuspended the transformed cells in 1 mL of SOC medium, regenerated them for 1 hour at 30 °C with shaking followed by overnight incubation on the bench, and then plated them on 100 μg/mL spectinomycin LB agar plates (100 μL of transformed cells per plate). We plated several microliters of diluted culture separately to determine the number of colony-forming cells. We incubated plates overnight at 37 °C and produced ∼2 × 10^6^ colonies in total. All randomly-picked colonies (over 50) failed to grow on ampicillin, suggesting that overnight incubation at 37 °C is sufficient to cure the majority of cells of the Tn7 helper plasmids. Similarly, we observed no growth on kanamycin, showing that the Tn7 integration plasmids were no longer present. We further validated the chromosomal incorporation of the barcode-carrying cassettes by colony PCR with a pair of primers directed to the Tn7 integration site. All randomly-picked colonies (over 50) were positive for the chromosomal integration. We scraped all the colonies from the plates, pooled them together, thoroughly mixed them, aliquoted them with 15% glycerol, and stored them at −80 °C.

### Deep sequencing of the gBlocks library, the Tn7 integration plasmid library, and the naïve barcoded *E. coli* population

Sample preparation for deep sequencing involved four steps. First, we amplified the barcode-carrying cassettes with Illumina adaptor primers (forward overhang: 5’ TCGTCGGCAGCGTCAGATGTGTATAAGAGACAG; reverse overhang: 5’ GTCTCGTGGGCTCGGAGATGTGTATAAGAGACAG). We used gBlocks and the Tn7 integration plasmid library directly as DNA templates in the PCR reaction. In the case of the naïve barcoded *E. coli* population, we performed the PCR reaction either directly on the cells or, alternatively, on the genomic DNA extracted from the barcoded population with Nucleospin Microbial DNA prep kit (Machary-Nagel). Second, we separated the PCR product on 1% agarose gel, excised it, and purified it using NucleoSpin Gel extraction kit (Machary-Nagel). Third, we subjected the gel-purified product of the first PCR reaction to a second PCR reaction using a pair of index primers from Nextera XT DNA library preparation kit (Illumina). Fourth, we purified the product of the second PCR reaction with Agencourt AMPure XP PCR purification kit (Beckman Coulter). We performed sequencing on MiSeq or NextSeq platforms (Illumina).

### Laboratory evolution

We subjected the naïve barcoded *E. coli* population to laboratory evolution via serial passaging under five distinct growth conditions: ‘low’ chloramphenicol (CMP) (1-3 μg/mL), ‘ultra-low’ CMP (0.1 μg/mL), ‘low’ trimethoprim (TMP) (0.1-1.2 μg/mL), ‘ultra-low’ TMP (0.01-0.1 μg/ml), and ‘no drug’ (**Fig. S3A,B**). We grew cells at 37 °C in a 96-well microtiter plate (500 μL per well) in supplemented M9 medium (0.2% glucose, 1mM MgSO4, 0.1% casamino acids, 0.5 mg/ml thiamin). Between passages, we grew cultures for 8-9 hours (during the day) or 10-12 hours (during the night). We used saturated culture from a previous passage to inoculate a fresh plate by 1:100 dilution (5 μL of saturated cultured into 500 μL of fresh medium). Overall, we performed 70 passages. We measured optical density (OD) of the cultures at 600 nm at the end of each passage. We convert the raw OD measurements to an estimated number of cells (**Fig. S3C,D**) as follows: we first multiply by 10 to correct for dilution of the measured culture, then multiply by 2 to standardize the OD to 1 cm path length, then multiply by cell density 10^8^ cells/mL/OD, and finally multiply by 0.2 mL as the volume of the well. We stored every second passage at −80 °C after addition of 15% glycerol.

### Fitness measurements of the evolving barcoded *E. coli* populations

We estimated fitness of the evolving populations by calculating the change in IC50 of chloramphenicol or trimethoprim. To this end, we sampled cells from three populations evolved in ‘low’ CMP (wells A1, B1, and C1), three populations evolved in ‘ultra-low’ CMP (wells A3, B3, and C3), and two populations evolved without antibiotics (wells E9 and F9) at passages 0, 8, 10, 12, 20, 30, and 70. We diluted these samples 1:100 into growth medium supplemented with 0, 1, 2, 4, 8, or 16 μg/mL of chloramphenicol, and followed their growth by OD measurements at 600 nm (**Figs. S2A, S4A**). We calculated the area under these growth curves over the time of growth (**Fig. S4B**) and normalized the area values so that they equaled 1 at zero antibiotic (**Fig. S4C**). We determined the IC50 by calculating the concentration of antibiotic at which growth (defined as normalized area under the growth curve) was reduced by 50% relative to zero antibiotic (**Fig. S4C**). We inferred the IC50 concentration by interpolating the area vs. drug concentration curves. We similarly obtained IC50 values for two populations evolved in ‘low’ TMP (wells A5 andB5), two populations evolved in ‘ ultra-low’ TMP (wells G7 and H7), and two populations evolved without antibiotics (wells E9 and F9), all at the same time points as for CMP. We used TMP concentrations 0, 0.5, 1, 2, 5, 10, and 20 μg/mL.

### Deep sequencing of the evolving barcoded *E. coli* populations

We sequenced barcodes at 16 time points over the evolution experiment for the same 12 independent populations used for the fitness measurements: three populations in ‘low’ CMP (wells A1, B1, and C1), three populations in ‘ultra-low’ CMP (wells A3, B3, and C3), two populations in ‘low’ TMP (wells A5 and B5), two populations in ‘ultra-low’ TMP (wells G7 and H7), and two populations in no antibiotics (wells E9 and F9). We first amplified these 192 bacterial cultures with Illumina adaptor primers (15 μL of defrosted cells from each culture). We performed the second PCR in two groups, each with 96 unique Nextera XT primers. We then pooled together 96 PCR reactions from each group, spiked them with 30% of PhiX DNA, and sequenced them on a Nextseq High Output 75 platform. The sequencing protocol commenced with 9 dark cycles to account for the sequence redundancy preceding the barcode area. For one sample – ‘ultra-low’ CMP, replicate 2 (well A3) at passage 54 – all PCR reactions failed, and thus we excluded this sample from all further analysis.

### Analysis of sequencing data

First, we exclude sequencing samples that report fewer than 10^6^ reads; this affects one of the samples from the initial population and six of the samples from the evolving populations (**Table S1**). All remaining samples have between 10^6^ and 7 × 10^6^ reads. Next, we exclude all reads with minimum base quality score less than 10 (Phred scale), which affects 0.02-0.05% of reads (**Table S1**). To identify barcodes, we first align each read to the reference sequence for the barcode cassette. We allow up to three mismatches or one indel with respect to the reference; we also require that the read overlap the barcode by at least 10 nt. With these criteria we identify barcodes on more than 95% of reads in almost all samples (**Table S1**). For each read with a valid alignment, we extract a barcode as the sequence aligning to the variable region in the reference.

To correct for sequencing errors in the raw barcodes, we use the bartender package^45^ on default settings to cluster together barcodes with similar sequences. In general, this method assumes that a low-frequency barcode differing at only one or two bases from a high-frequency barcode is the result of a sequencing error, so that the low-frequency barcode is merged into the high-frequency one. This produces a set of putatively true barcodes for the sample.

To ensure that we identify true barcodes consistently across samples, we first pool raw barcodes and perform clustering on these pooled samples. We pool barcodes both across time points for each population (to build trajectories of barcodes over time in each population) as well as across populations for each time point (to compare barcodes between populations). After clustering we disaggregate the true barcodes from the pooled data back into the individual samples, where we normalize them by the total number of reads in that sample. This yields a set of lineage frequencies {*x_k_*} (where the index *k* runs over all barcodes) for each population at each time point.

### Quantifying lineage diversity

The simplest way to quantify the diversity of lineages in a population is to count the number of unique barcodes observed at a particular time point (**Fig. 3A**). However, if lineages differ widely in frequency, then this measure may not be very informative and will suffer from significant sampling bias (since very low-frequency barcodes will be under-sampled). A more general approach is to define the *effective* number of lineages using the diversity index *q_D_* from ecology^34^. We construct this definition in analogy with the case where all lineages are at equal frequency, so that the number of lineages is simply the reciprocal of this frequency:

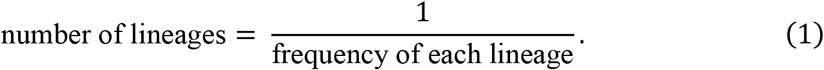

When lineages are not at equal frequencies, we replace the frequency in the denominator by a mean frequency over all lineages. Define the generalized mean (also known as the power mean) of a quantity *h*_k_, with normalized weights *p_k_* (Σ_*k*_ *p_k_* = 1) and parameter *q*:

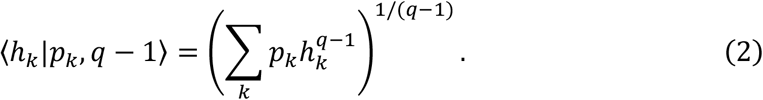

The parameter *q* controls how strongly the mean depends on very small or very large values of *h_k_*: lower values of *q* more strongly weigh low values of *h_k_*, while higher values of *q* weight high values of *h_k_* more strongly. When *q* = 2, Eq. 2 reduces to the ordinary arithmetic mean; when *q* = 1, it is equivalent to the geometric mean; and when *q* = 0, it is the harmonic mean.

We therefore define the effective number of lineages as the reciprocal of the generalized mean frequency over all lineages, weighing each frequency by itself^34,35^:

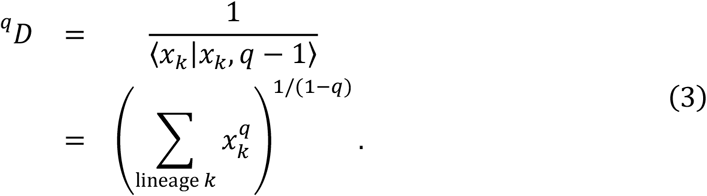

The diversity index is mathematically equivalent to the exponential of the Renyi entropy in physics^46^. Special values of the parameter *q* correspond to common ecological measures of diversity:

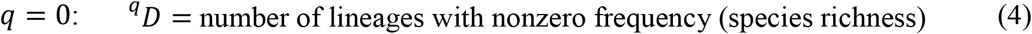

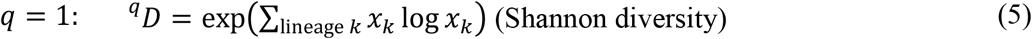

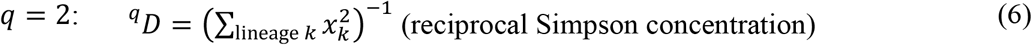

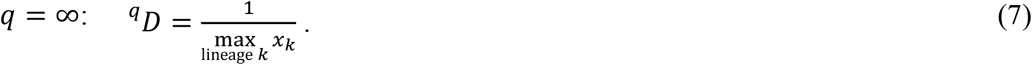

Note that the effective number of lineages according to Eq. 3 equals the actual number of lineages for any *q* if all lineages have equal frequency. **Figure 3** shows the diversity indices for *q* = 0, *q* = 1, and *q* = ∞ for each population over the course of the evolution experiment.

### Quantifying dissimilarity of lineage composition between populations

We can also use ecological diversity indices to quantitatively compare the lineage compositions of two or more populations. Let *M* be the number of populations we comparing (*M* ≥ 2), and let 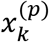 be the frequency of lineage *k* in population *p* (= 1, 2, …, *M*) at a particular time point. If we pool together all *M* populations, the frequency of lineage *k* is

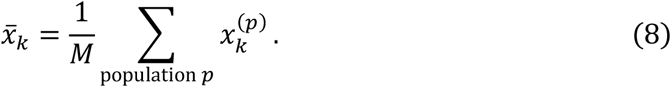

The total diversity of the pooled population (“gamma diversity”) is^34,35^:

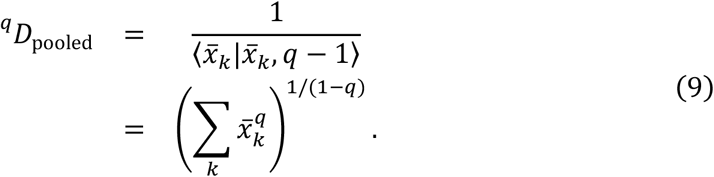

We can decompose this total diversity into two factors:

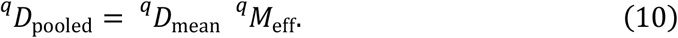

The first factor on the right-hand side of Eq. 10 is the mean diversity across all populations (“alpha diversity”):

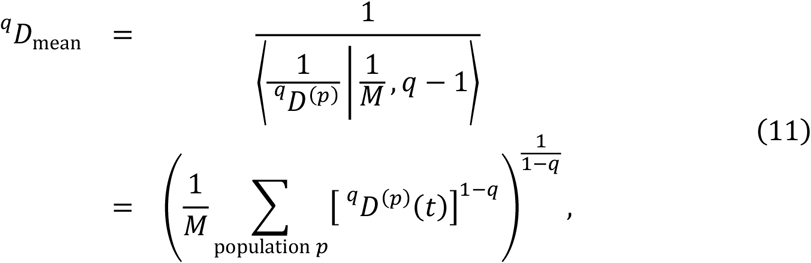

where ^*q*^*D*^(*p*)^ is the diversity of population *p* alone (Eq. 3). The second factor on the right-hand side of Eq. 10 is the effective number of distinct populations (“beta diversity”):

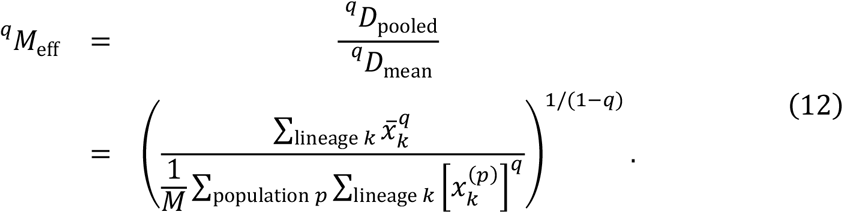

This quantity has a minimum value of 1 if the populations have identical lineages at identical frequencies, and a maximum value of *M* if none of the populations have any lineages in common. To simplify the interpretation of this quantity across cases where we may be comparing different numbers of populations (e.g., three replicates in ‘low’ CMP versus two replicates in ‘low’ TMP), we shift and rescale ^*q*^*M*_eff_ to obtain a measure of dissimilarity between populations that ranges from 0 to 1:

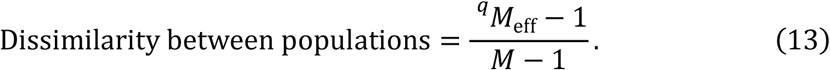

We plot this normalized quantity in all figures (**Figs. 4, S8, S9**).

In the case of *q* = 0, ^*q*^*M*_eff_ simply measures how many lineages are in common between the populations under comparison. Let *B*^(*p*)^ be the set of lineages with nonzero frequencies in population *p*, and let |*B^(p)^*| denote the number of lineages in this set. Then the effective number of distinct populations is

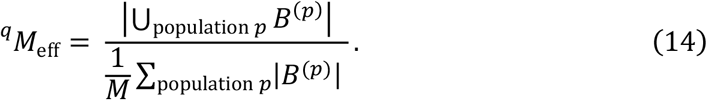

For two populations (*M* = 2), we can rewrite this as

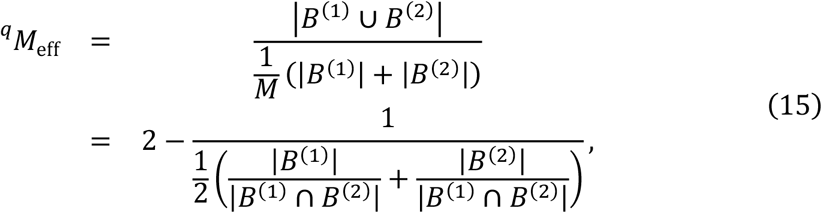

where we have invoked the inclusion-exclusion principle for sets: 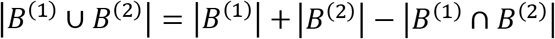. That is, ^*q*^*M*_eff_ equals two minus the harmonic mean of the fractions of overlapping lineages between the populations. For example, ^*q*^*M*_eff_ = 1.8 means that the two populations have 20% of their lineages in common.

In the case of *q* = 1, the effective number of lineages is the Shannon diversity (Eq. 5). Therefore the effective number of distinct populations is equivalent to the exponential of the Jensen-Shannon divergence between the populations:

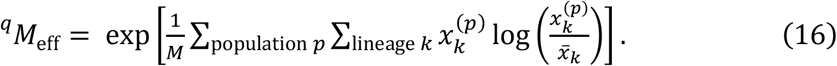

This is also equivalent to the weighted sum of the Kullback-Leibler divergences between each population and the pooled population.

In the case of *q* = ∞, the effective number of distinct populations depends only on the most abundant lineage in the pooled population and across all populations. That is,

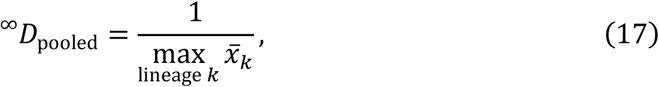

and

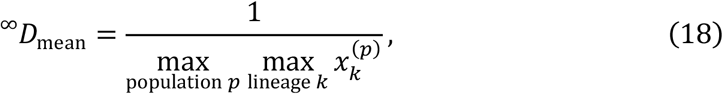

and so

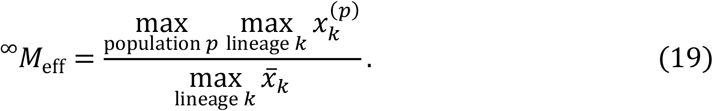

### Clustering lineage frequency trajectories

For each population, we exclude barcoded lineages that have zero detected frequency at more time points than a minimum number calculated as

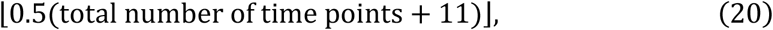

where ⌊ ⌋ is the floor function that rounds the argument down to the nearest integer. This ensures that all pairs of remaining lineages have at least 10 time points at which they both have nonzero frequency. This leaves between 310 and 695 lineages for each population. We cluster the frequency trajectories *x_k_*(*t*) for these lineages using the hierarchical clustering routine in SciPy^47^. The distance metric between two lineages *k*_1_ and *k*_2_ is

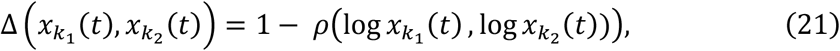

where *ρ*(log*x*_*k*_1__(*t*), log *x*_*k*_(*t*))) is the Pearson correlation coefficient between the logarithms of both frequency trajectories (excluding time points where either frequency is zero); **Fig. S13** shows matrices of all pairwise trajectory distances. We use the “average” linkage method (equivalent to unweighted pair group method with arithmetic mean, or UPGMA), which calculates the distance between two clusters as the arithmetic mean of the distances between all trajectories in both clusters. Other linkage methods produce qualitatively similar results. The hierarchical clustering results in dendrograms as shown in **Fig. S13**. Finally, we form flat clusters by setting thresholds on the dendrograms, which we manually choose for each population; these thresholds are shown on the dendrograms in **Fig. S13** and range from 0.35 to 0.65, which roughly mean that the correlation between trajectories within clusters is at least 0.35 or 0.65.

## Acknowledgments

We thank Lu Zhao, Jose Rojas Echenique, Sasha Levy, and Alberto Pascual Garcia for valuable discussions and advice, and Dan Tawfik and Amir Aharoni for insightful comments and help with preparation of the manuscript. MM was supported by an F32 fellowship from the National Institutes of Health (GM116217) and an Ambizione grant from the Swiss National Science Foundation (PZ00P3_180147). A.W.R.S. acknowledges support from the Canadian Natural Sciences and Engineering Research Council (NSERC RN000524). This work was supported by personal Israel Science Foundation grant 1630/15 to SB.

## Authors Contribution

S.B. conceived the study. J.L. and W.J. performed the laboratory experiments. M.M. analyzed sequence data and applied ecological tools. S.B., M.M., A.W.R.S., L.G., analyzed the data. S.B. and M.M. wrote the manuscript. All authors approved of the final manuscript.

## Supplementary Figures

**Figure S1:**
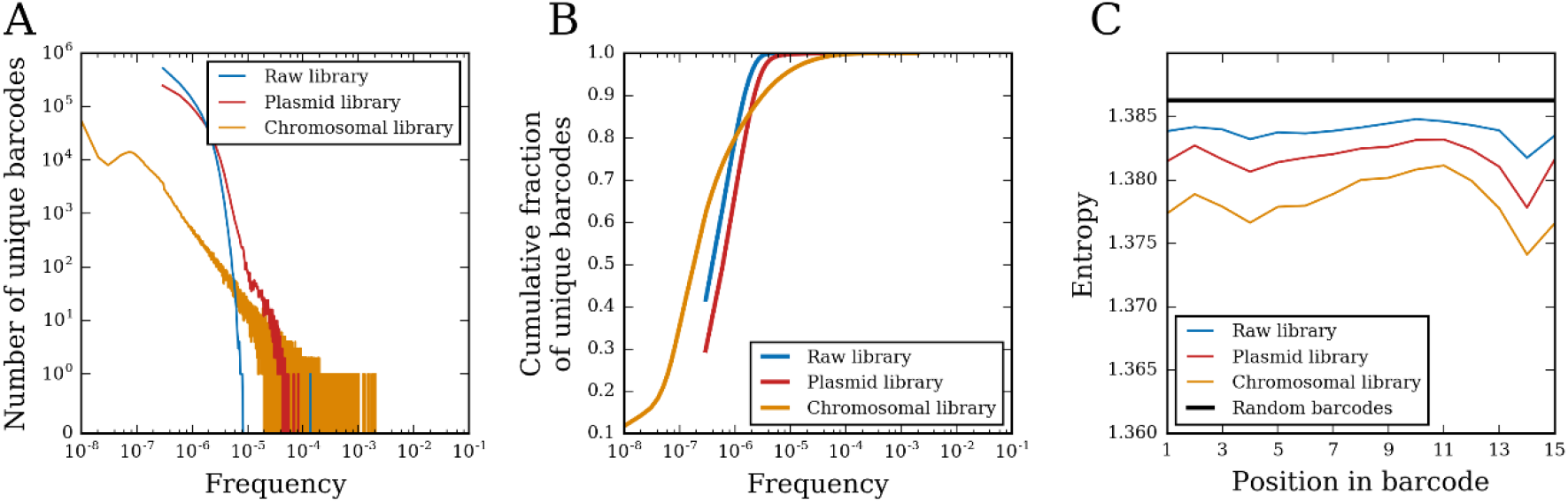
Statistics of the initial barcode libraries. **(A)** Distributions of barcode frequencies at different stages of library preparation. For a given frequency on the horizontal axis, the vertical axis shows the number of unique detected barcodes with that frequency. ‘Raw library’ (blue): NextSeq Illumina sequencing of the barcode library as synthesized by IDT (prior to plasmid library creation). ‘Plasmid library’ (red): MiSeq Illumina sequencing of barcodes incorporated into the Tn7 integration plasmid library. “Chromosomal library” (orange): NextSeq Illumina sequencing of the barcode library integrated into *E. coli* chromosomes and generated by PCR performed on chromosomal DNA pooled from five independent extractions. **(B)** Same as panel (A), but showing the cumulative distributions of frequencies. **(C)** Shannon entropy of nucleotides at each position in the 15 nt barcode for the same libraries in panel (A). The horizontal black line marks the entropy (ln 4 ≈ 1.386) for a truly random library of barcodes, where all nucleotides are equally abundant at each position.

**Figure S2:**
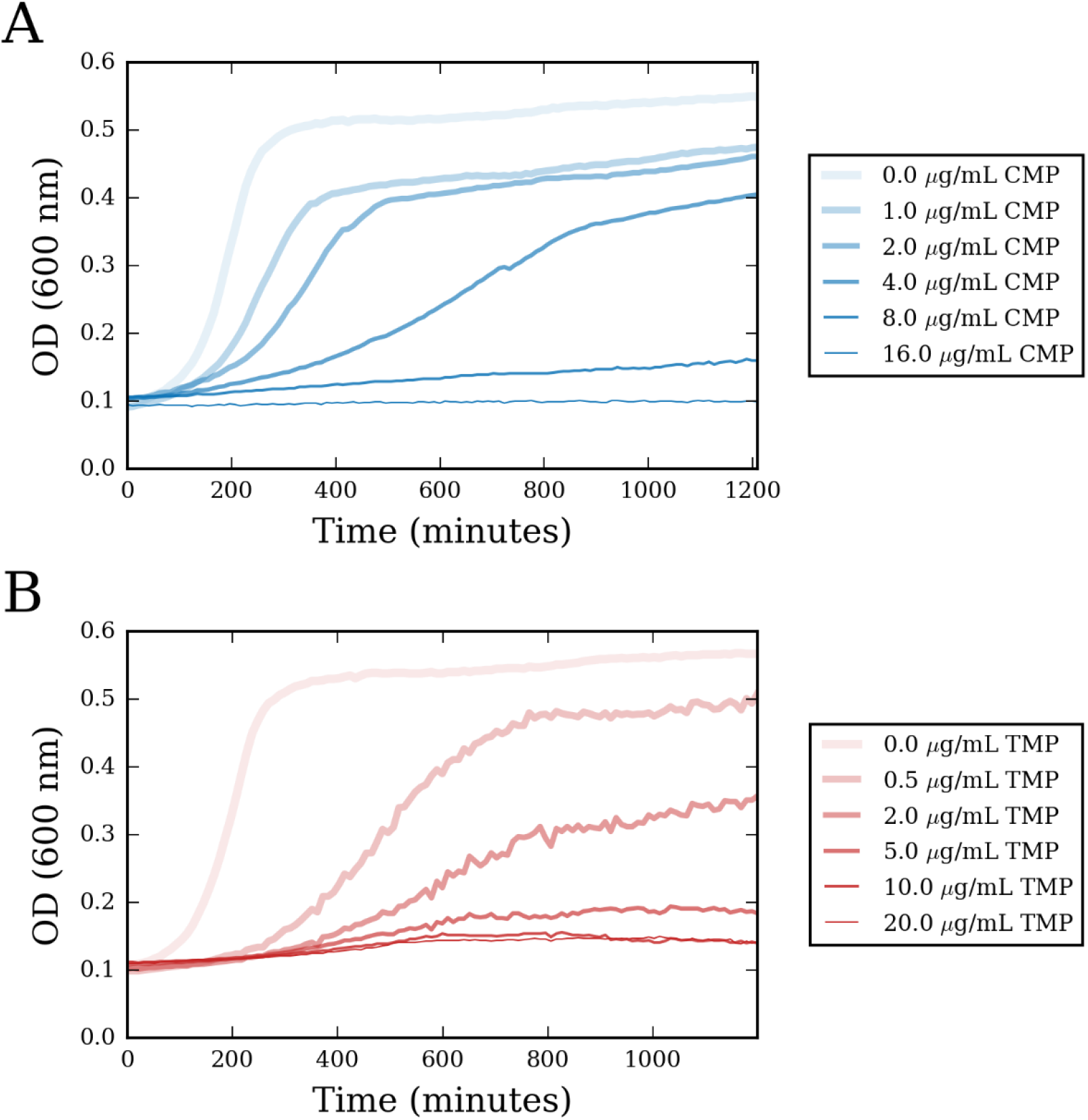
MIC values of the naïve barcoded population. We grew the naïve barcoded population for 20 hours in the presence of **(A)** 0-16 μg/ml of chloramphenicol (CMP) or **(B)** 0-20 μg/ml of trimethoprim (TMP). We defined the MIC for each drug as the lowest concentration of antibiotic at which we observed no growth.

**Figure S3:**
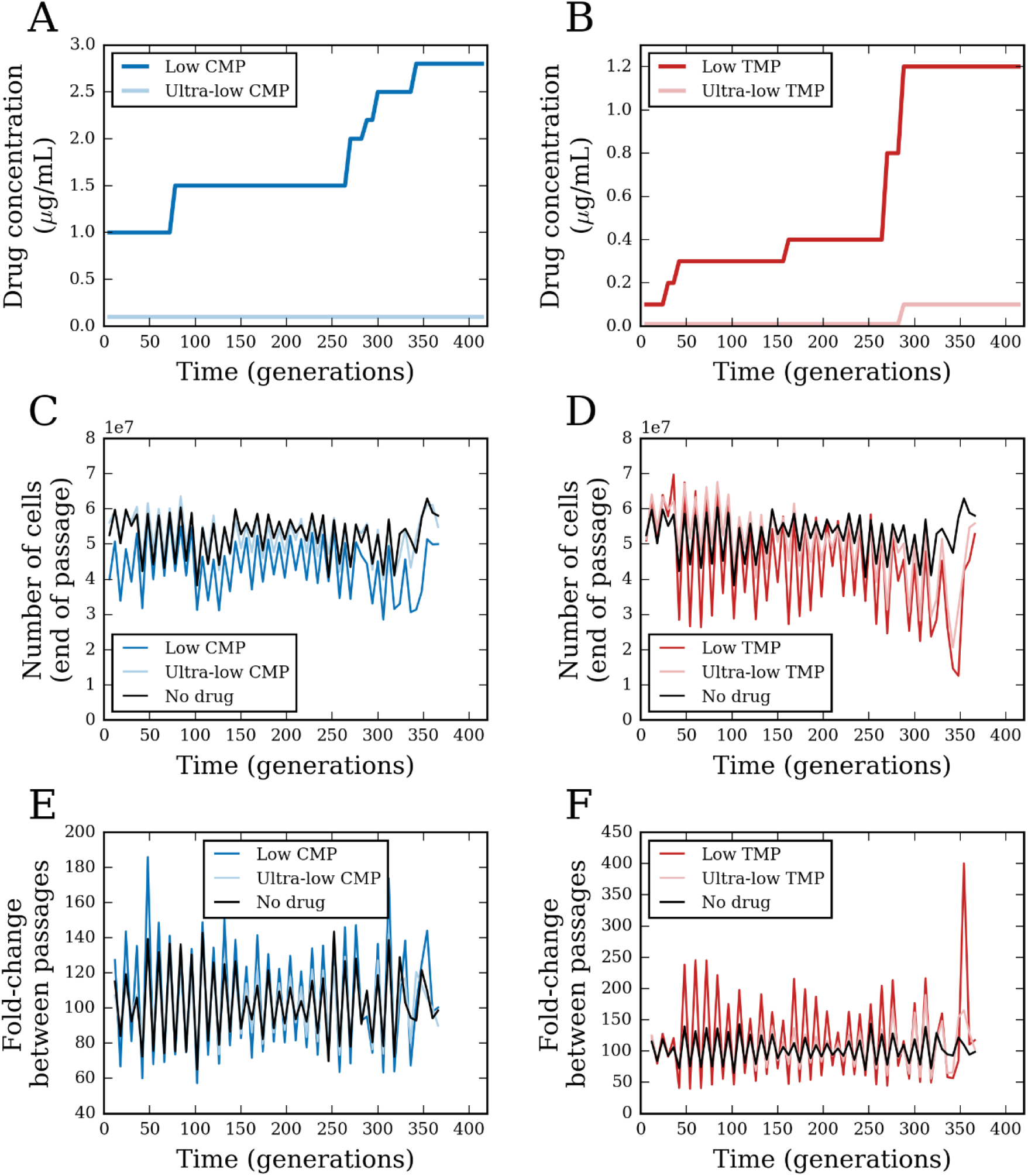
Drug concentrations and population growth over evolution experiment. **(A)** Trajectories of ‘low’ and ‘ultra-low’ chloramphenicol (CMP) concentrations over time of the evolution experiment. **(B)** Same as (A) but for trimethoprim (TMP) conditions. **(C)** Approximate number of cells at the end of each passage for ‘low’ and ‘ultra-low’ CMP conditions, along with the populations evolved without drug. Lines are averages over all 14 replicate populations for each condition. **(D)** Same as (C) but for TMP conditions. **(E)** Same as (C) but showing the fold-change of population size during each passage on the vertical axis. **(F)** Same as (E) but for TMP conditions. Periodic oscillations in cell numbers and yields result from the fact that cultures were propagated in two intermittent growth regimes: 9 hours during the day, followed by 12 hours during the night (see **Methods**).

**Figure S4:**
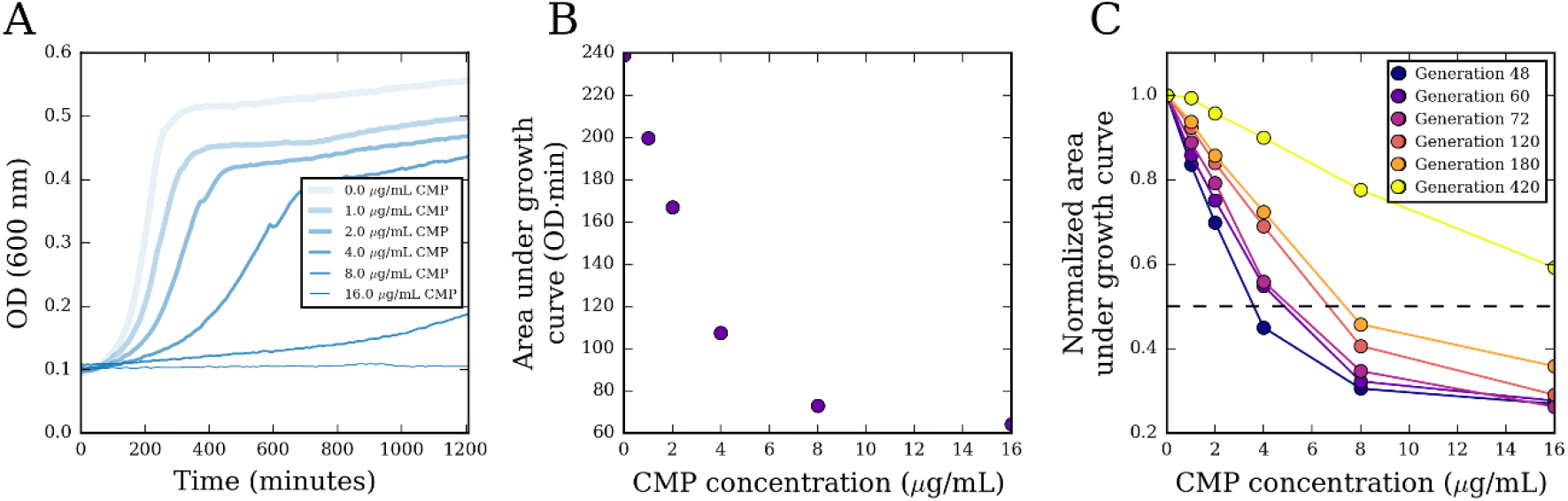
Example of IC50 calculation. **(A)** Growth curves of cells from a barcoded population evolving in ‘low’ chloramphenicol (CMP) at generation 120 (passage 8), measured with different concentrations of CMP. **(B)** For each growth curve in (A), we calculate the area under it and plot the area as a function of CMP concentration. **(C)** We similarly calculate growth curve areas for generations 48, 60, 72, 180, and 420 and normalize them by the area of the growth curve with zero CMP. The IC50 is then defined as the antibiotic concentration leading to 50% of growth relative to growth at zero CMP.

**Figure S5:**
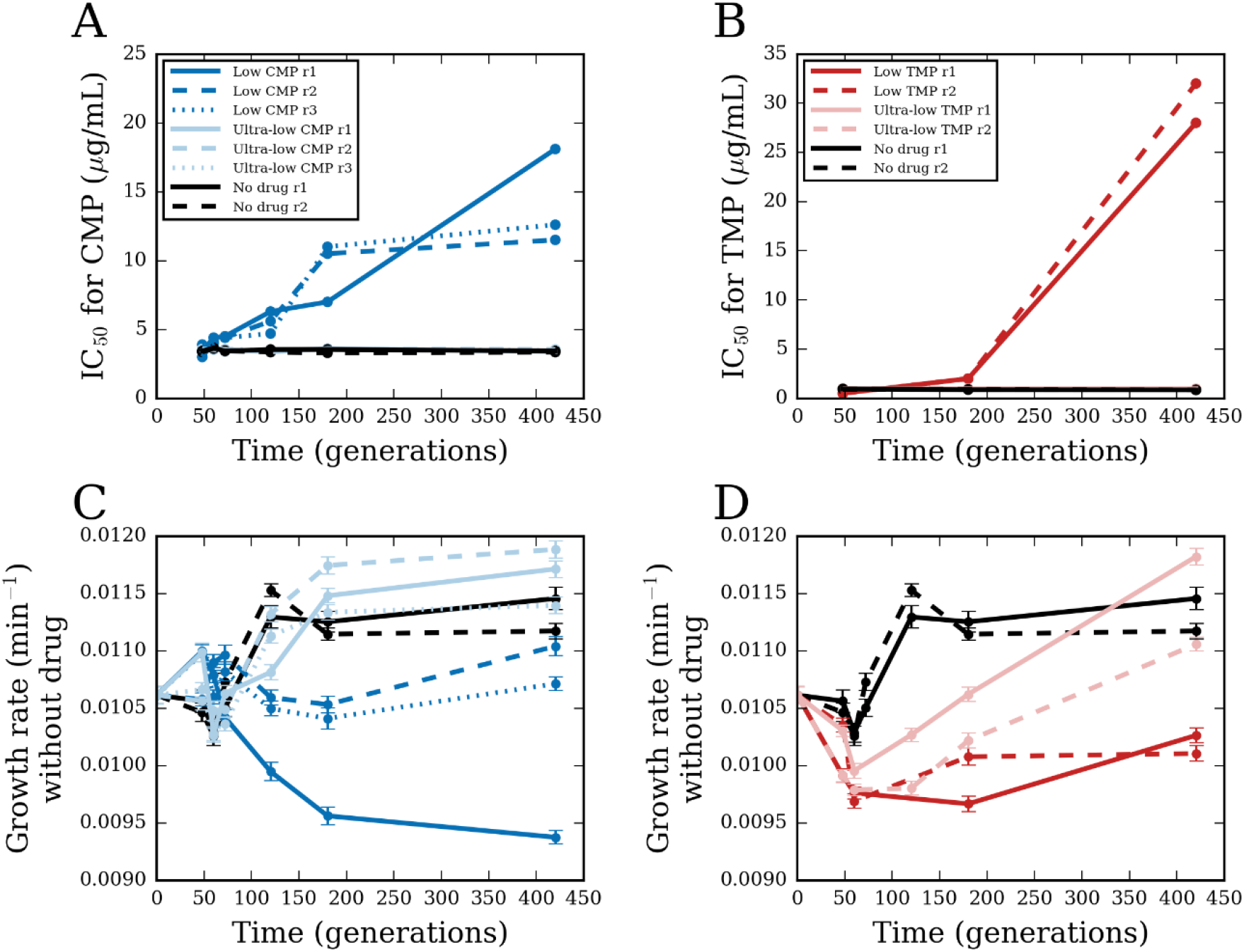
Fitness of the evolved populations. **(A)** Chloramphenicol (CMP) concentration inhibiting 50% of growth (IC50) of the barcoded populations evolving in ‘low’ and ‘ultra-low’ CMP as well as without drug. **(B)** Same as (A) but for trimethoprim (TMP). **(C)** Growth rate, measured in the absence of drug, of barcoded populations evolving in ‘low’ and ‘ultra-low’ CMP as well as without drug. Points represent the mean and error bars represent standard deviation over replicate measurements. **(D)** Same as (C) but for populations evolved in ‘low’ and ‘ultra-low’ TMP.

**Figure S6:**
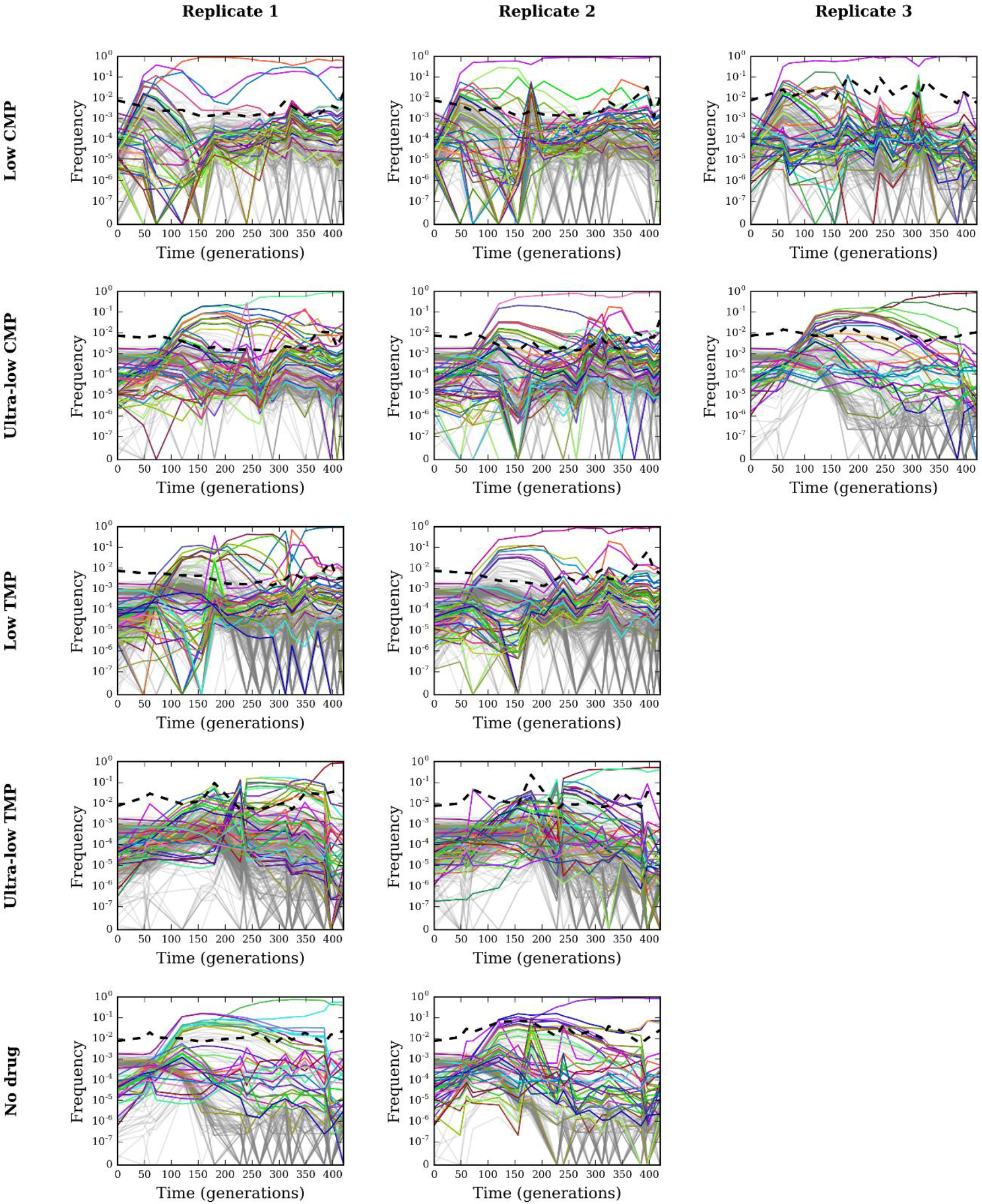
Trajectories of barcoded lineage frequencies. Frequency trajectories for barcodes with average frequency greater than 10^−4^. Each row corresponds to a different condition and each column corresponds to a different replicate. Lineages that rank in the top 10 (according to average frequency) in any population are colored in all panels (**Table S2**); the colors are consistent across panels and match **Fig. 2**. Lineages that do not rank in the top 10 for any population are in gray and made transparent to show their density. The dashed black line shows the frequency of reads without identified barcodes.

**Figure S7:**
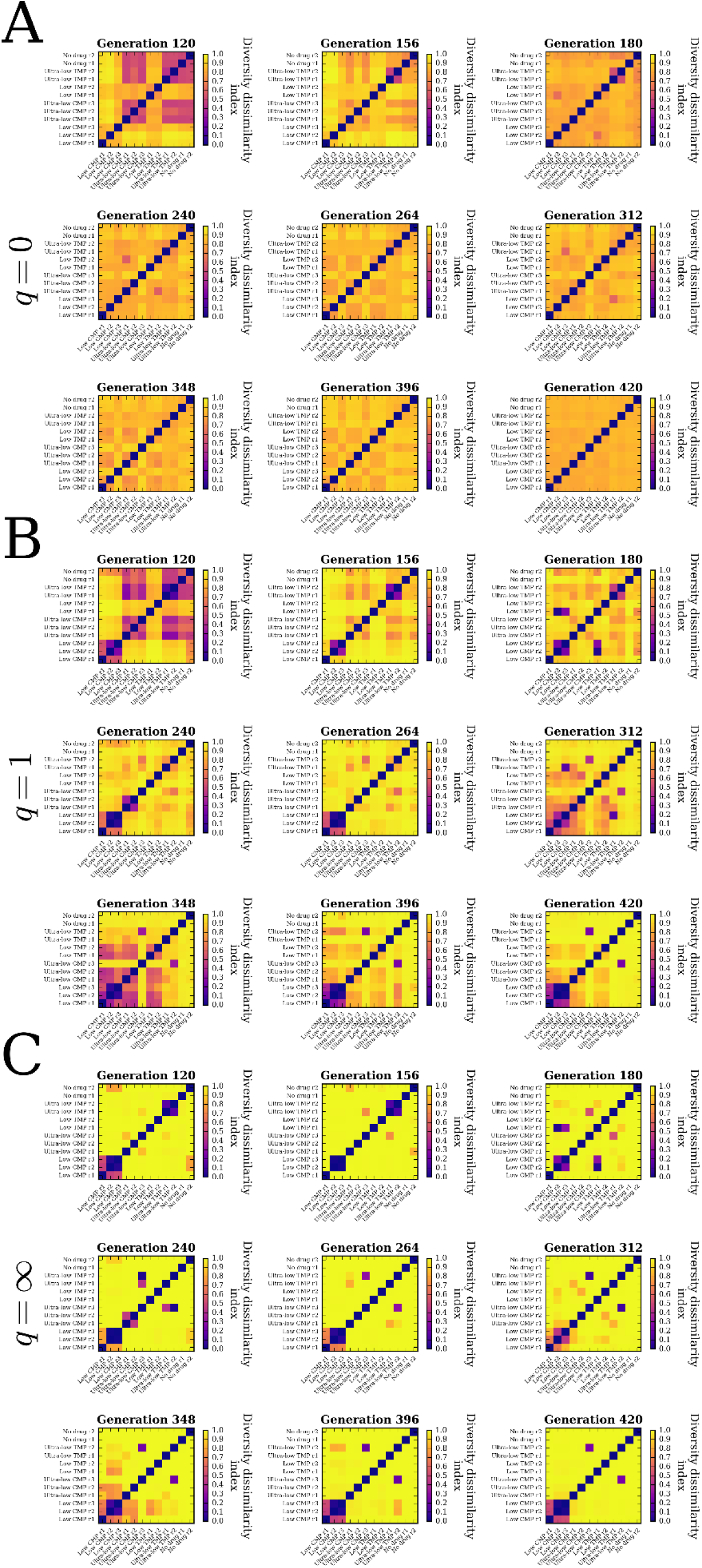
Dissimilarity of lineage frequencies between populations. Each panel shows the diversity dissimilarity index (**Eq. 2, Methods**) between all pairs of populations at a particular time point: **(A)** *q* = 0, (B) *q* = 1, **(C)** *q* = ∞.

**Figure S8:**
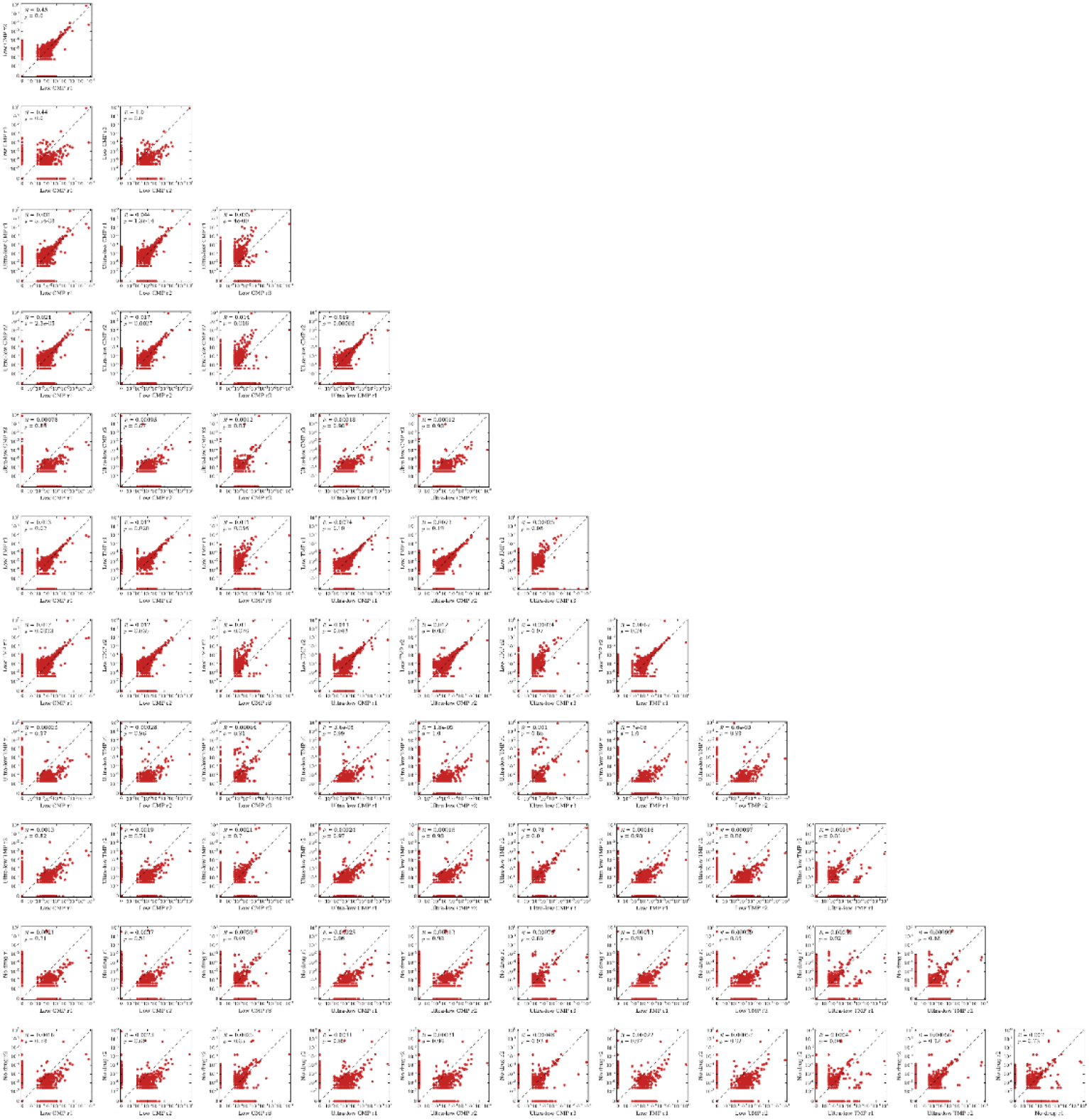
Correlation of lineage frequencies between populations. Scatter plots comparing lineage frequencies at the end of the experiment between each pair of populations. Each point represents a unique barcoded lineage; the dashed line is the line of identity.

**Figure S9:**
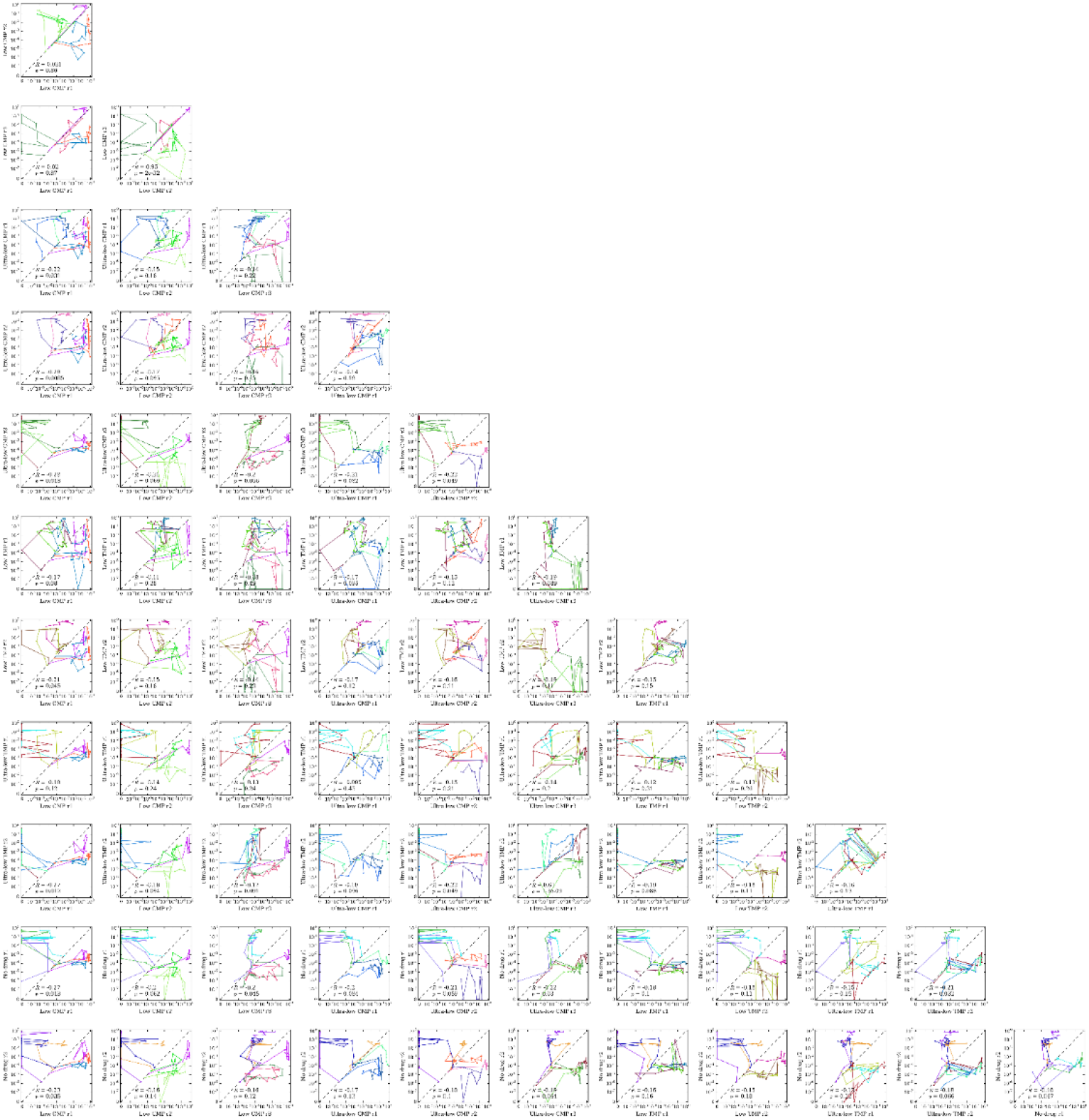
Correlation of top lineage trajectories between populations. Plots show traces of the top three (by average frequency) lineages over time between each pair of populations. Colors of lineages are consistent across panels and match **Fig. 2**. The dashed line is the line of identity.

**Figure S10:**
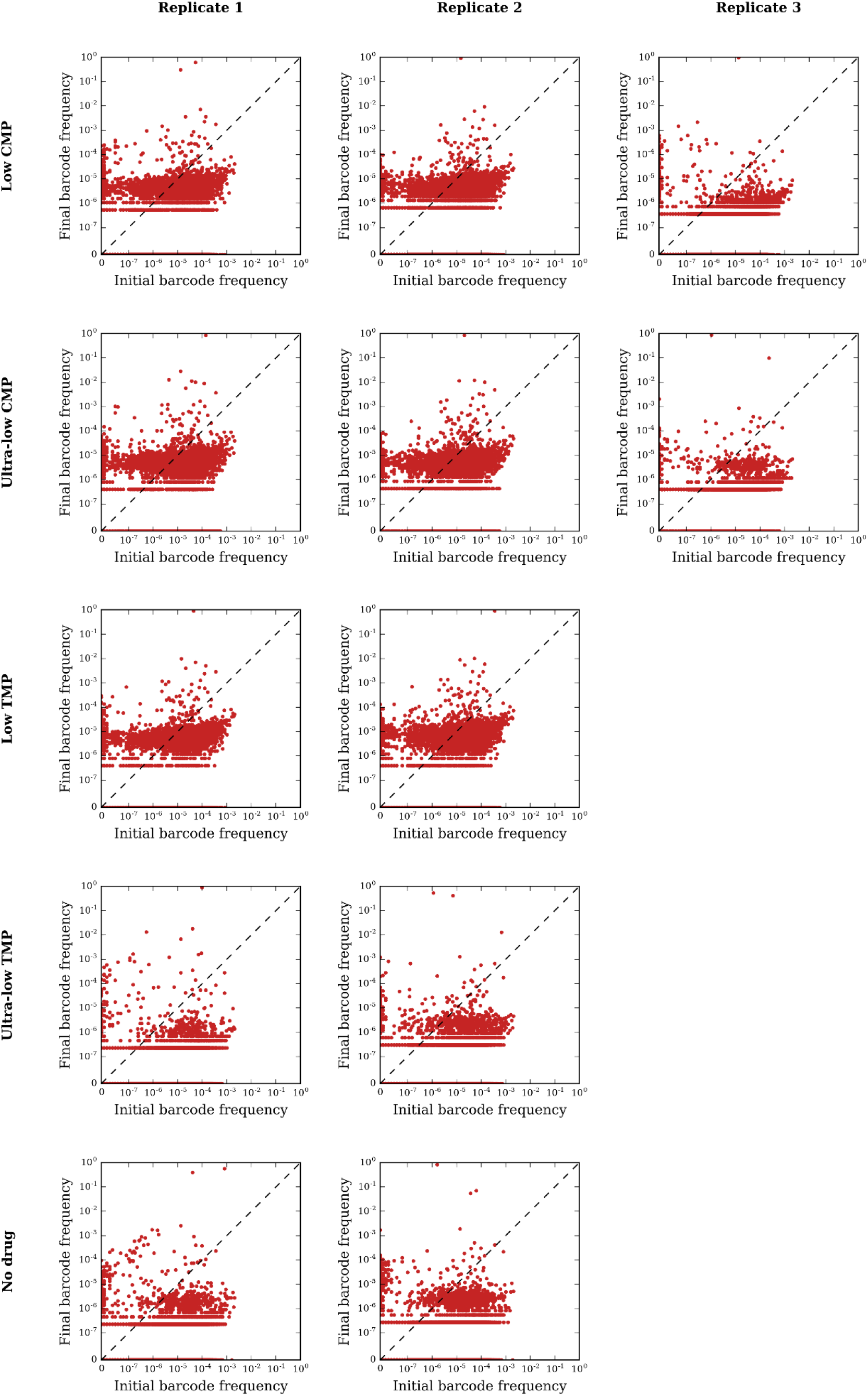
Correlation of lineage frequencies before and after evolution. Scatter plot comparing lineage frequencies at the final time point (vertical axis) with frequencies at the beginning of the experiment (horizontal axis). Each point represents a single lineage; the dashed line is the line of identity. Each row corresponds to a different condition and each column corresponds to a different replicate.

**Figure S11:**
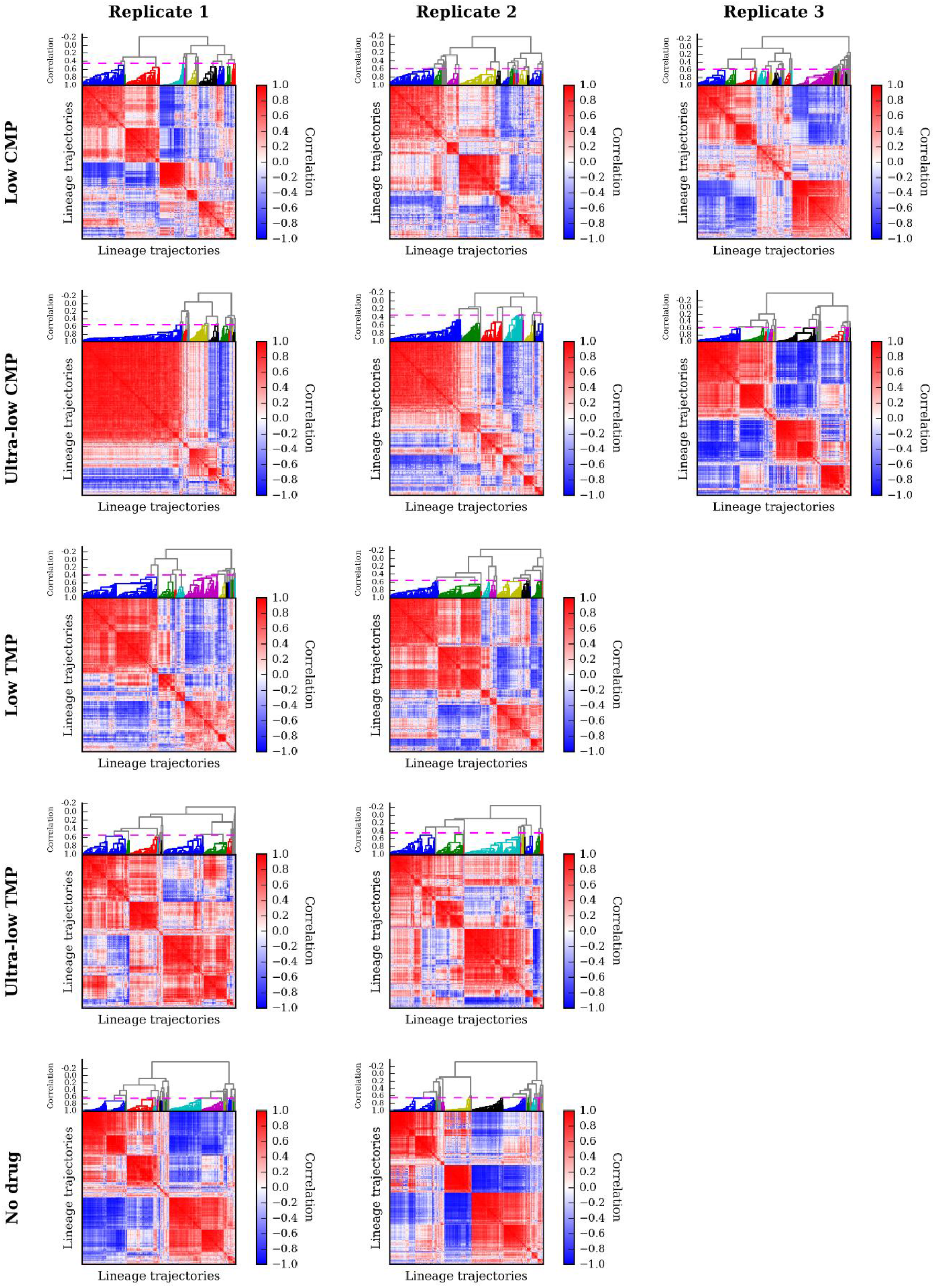
Summary of trajectory clustering. In each panel we show the matrix of Pearson correlation coefficients between all pairs of trajectories used for hierarchical clustering and the resulting dendrogram (Methods). The horizontal dashed line marks where we cut off the dendrogram to form flat clusters used for further analysis. Each row corresponds to a different condition and each column corresponds to a different replicate.

**Figure S12:**
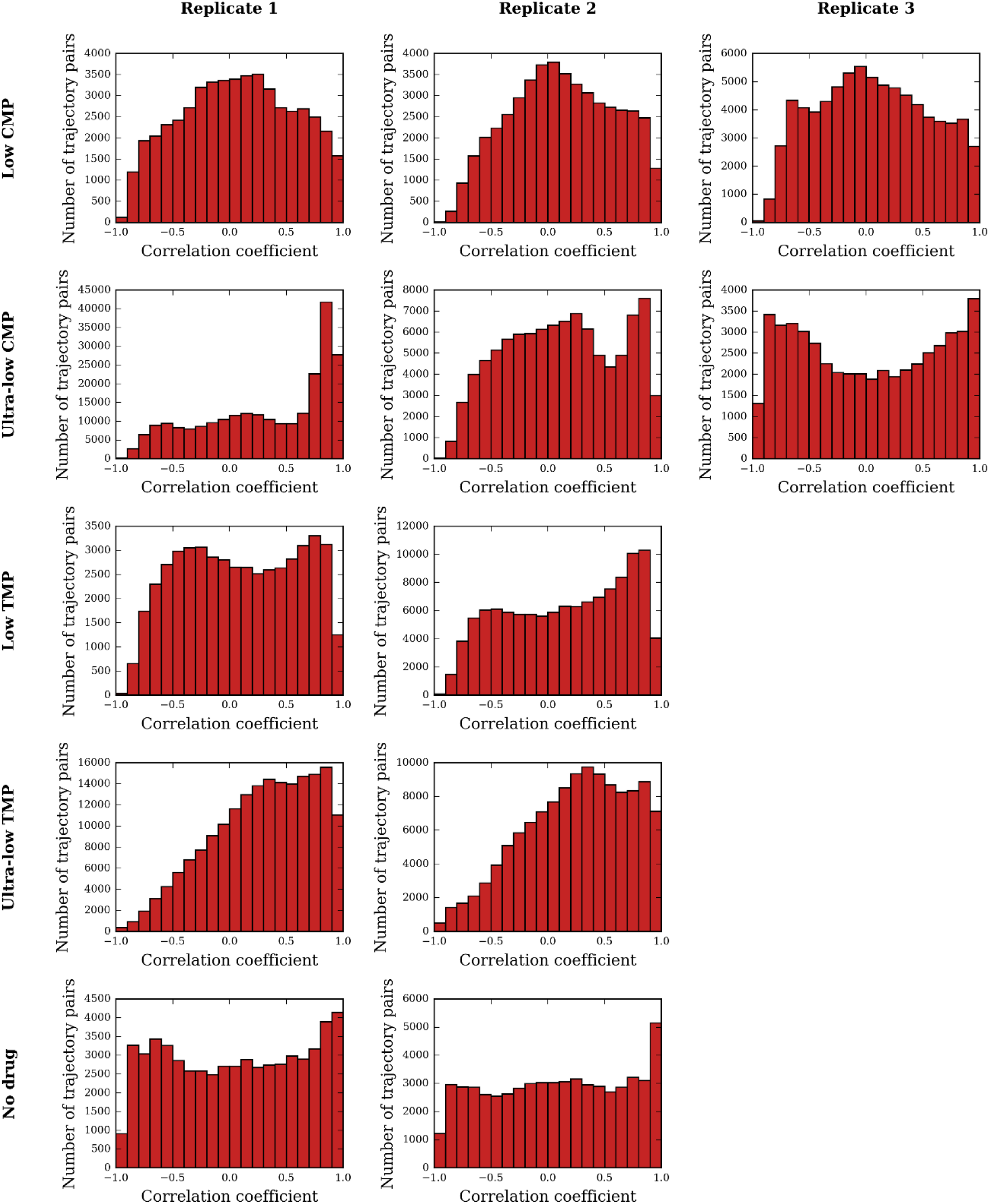
Distributions of trajectory correlations. In each panel we show the histogram of Pearson correlation coefficients between all pairs of lineage trajectories used for hierarchical clustering population (see **Methods**). Each column corresponds to a different condition and each row corresponds to a different replicate.

**Figure S13:**
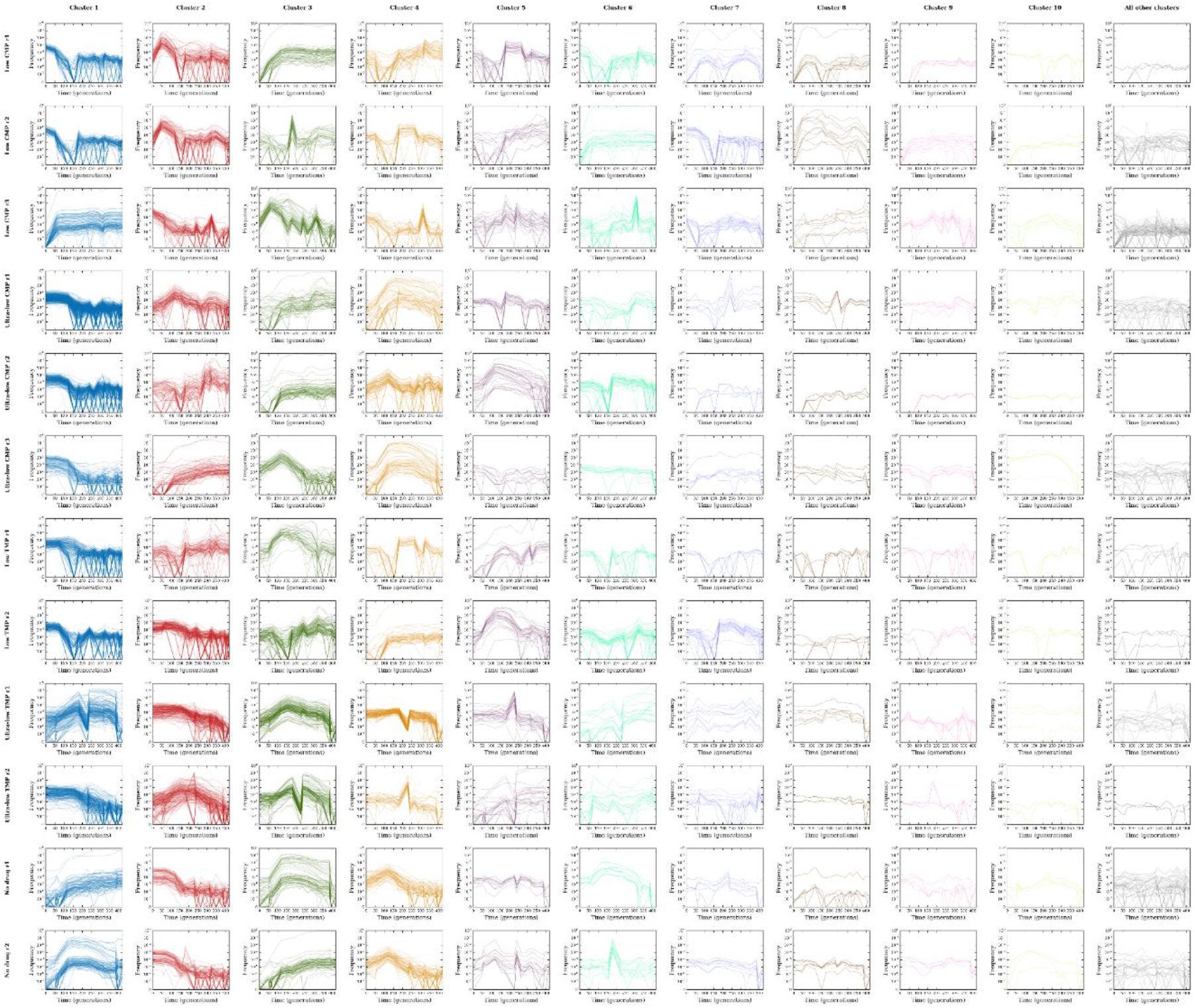
Clusters of lineage trajectories. Each panel shows a set of lineage trajectories that clustered together in our analysis. Each row corresponds to a single population. The first 10 columns show the top 10 clusters in decreasing order of cluster size (number of trajectories), while the last column shows trajectories from all other clusters.

**Figure S14:**
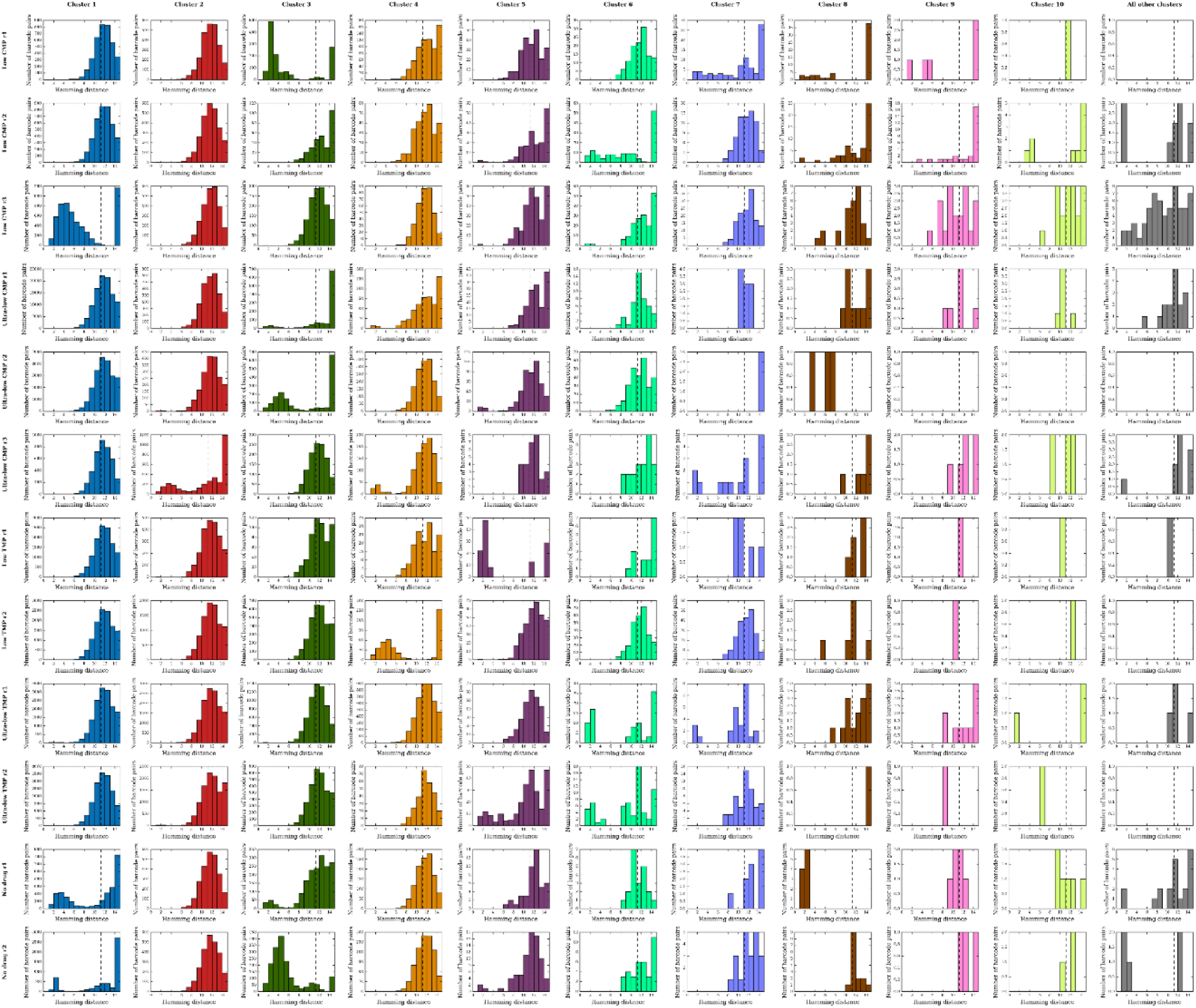
Hamming distances among sequences within trajectory clusters. Each panel shows the histogram of Hamming distances between all pairs of barcode sequences whose trajectories clustered together in our analysis. The first 10 columns show the top 10 clusters in decreasing order of cluster size (number of trajectories), while the last column shows trajectories from all other clusters. The vertical dashed line marks the expected Hamming distance between two random barcode sequences (11.25). Empty histograms correspond to clusters with only one trajectory.

